# Variation in bud phenology, frost tolerance and non-structural carbohydrates among white spruce seed sources on climate-contrasted test sites: implications for assisted migration

**DOI:** 10.64898/2025.12.10.693555

**Authors:** Chafik Analy, Lahcen Benomar, Martin Perron, Julie Godbout, Jean Beaulieu, Yves Bergeron, Jean Bousquet, Mebarek Lamara

**Affiliations:** Institut de recherche sur les forêts, Université du Québec en Abitibi-Témiscamingue, Rouyn-Noranda, QC, Canada; Ontario Forest Research Institute, Ontario Ministry of Natural Resources, 1235 Queen Street East, Sault Ste. Marie, ON, Canada; Direction de la recherche forestière, Ministère des Ressources naturelles et des forêts, Québec, QC, Canada; Centre d’étude de la forêt et Institut de Biologie intégrative et des systèmes, Faculté de foresterie, de géographie et de géomatique, Université Laval, Québec, QC, Canada; Département des Sciences biologiques, Université du Québec à Montréal, Montréal, QC, Canada

**Keywords:** bud set phenology, *Picea glauca*, assisted migration, cold hardiness, functional traits, growth, non-structural carbohydrates

## Abstract

To assess the potential impacts of climate change on white spruce local adaptation, six seed sources were evaluated seven years after plantation on two test sites with contrasting growing conditions and latitude in Quebec. Bud set, frost tolerance and analysis of non-structural carbohydrates (NSC) content showed important effects of test sites, seed sources, and their interactions on bud set phenology and growth. The average bud set initiation occurred one week later in the southern site compared to the northern site, whereas the late stage of bud set was similar between sites. Significant differences were observed between seed sources for some phenological stages. Frost tolerance was significantly lower in the southern site and below -12 °C for all seed sources sampled at the beginning of October. The trend in fructose and glucose content was opposite between sites in September. It decreased from September to October in the southern site and increased in the northern site, while sucrose content showed an opposite pattern, with the southwestern seed source harboring the lowest sucrose content. NSC content was also correlated to frost tolerance. Our study highlighted the crucial role of NSC in cold hardiness to early fall frosts and local adaptation. Implications for assisted migration are discussed.

## INTRODUCTION

During past periods of climate change, natural forest species responded individually with extensive geographic migrations to and from glacial refugia (Petit et al., 2008; Gonzales et al., 2009; Jaramillo-Correa et al., 2009). Recent research showed that many tree species are already undergoing distributional changes in response to the current global warming, highlighting the poleward and upper elevation shifts of some species (Chen et al., 2011; Aidenapol et al., 2015; Wu et al., 2015), and others moving in longitude following moisture availability and temperature changes (Godbout et al., 2012; Fei et al., 2017).

In Canada, it is predicted that the boreal forest will experience greater environmental variation in temperature and precipitation than the global averages (Adam-Poupart et al., 2014). In the eastern part of the boreal forest, the temperature is expected to increase by up to 6 °C in the horizon of 2100 under the RCP8.5, the worst-case scenario, coupled with an overall increase in precipitation (IPCC, 2021). For the boreal forest of Quebec (Eastern Canada), the Ouranos Consortium on Regional Climatology and Adaptation to Climate Change has predicted similar increases for the years 2071-2100, where mean annual temperatures could rise between 4 °C and 7 °C as one moves northward in Quebec. However, the climate change underway is shifting the environmental conditions too fast for trees to follow by migration alone their optimal growing conditions (Aitken et al., 2008; Alberto et al., 2013; Sittaro et al., 2017). Already, signatures of local genetic maladaptation in white spruce (*Picea glauca* [Moench] Voss) have been recorded with regards to warming temperatures (Andalo et al., 2005; Benomar et al., 2022). Despite the obvious warming trend in mean temperature, climate change is also expected to increase the frequency and severity of extreme events, which have a high potential to have a major impact on vegetation communities (Lloret et al., 2012; Marquis et al., 2022).

Assisted population migration, also called assisted gene flow, has been proposed as a proactive approach to help forest species adapt to global change. It has also been suggested as a strategy to mitigate the effect of climate change on forests (Pedlar et al., 2012; Aitken & Bemmels, 2016). One way to apply these principles is by developing transfer models that rely on the calculation of the maximum transfer distance of seed sources, which is generally within the natural range of the species distribution, using growth response and bioclimatic predictors (Beaulieu & Rainville, 2005). However, when moving local seed sources to colder environments, assisted migration may carry the risk of frost damage if seed sources are moved too far, or if expected warmer climate conditions have not yet been established (Sebastian-Azcona et al., 2018; Benomar et al., 2022). Despite the warming trend observed in the recent decades in northern mid-latitudes, frost damage has still occurred in natural populations, particularly early in the growing season. For example, severe frost events were recorded in spring 2007 in Eastern North America (Gu et al., 2008; Man et al., 2009), and again in 2021 following an early warm spring, when a late spring cold spell caused severe damages to both local populations and migrated seed sources (Benomar et al., 2022). Therefore, frost damage and growth limitations became a serious risk for newly implemented populations.

Differences among seed sources in bud set and the onset of cold hardiness during late summer and in the fall have also been reported to be greater than those for bud flush and the release of cold hardiness in spring (e.g., Li et al., 1993). Thus, latitudinal transfers might have a greater impact on susceptibility to early fall frost occurrences (Aitken & Hannerz, 2001). Several studies have highlighted the necessity of including ecophysiological responses and non-structural carbohydrates metabolism variables in seed transfer models to reduce frost damage risks (Villeneuve et al., 2016; Marquis et al., 2020; Benomar et al., 2020, 2022). Such models should include extreme weather events, cold hardiness chronology, non-structural carbohydrates dynamics of seed sources, and ecological conditions of test sites at the end of the growing season. Knowing that cold hardiness heavily relies on bud set phenology and the dormancy depth, and that variation for this trait is usually the main factor affecting the length of the growing season (Li et al., 1993; Sebastian-Azcona et al., 2018), the trade-off between growth and cold hardiness is largely driven by how long trees extend their growing season during the fall (Howe et al., 2003).

In this study, we used six white spruce seed sources (collected in seed orchards) originating from different bioclimatic domains in the province of Quebec in Canada (Benomar et al., 2018), and local population controls to assess the importance of adaptive traits that could impact tree productivity in two sites presenting contrasted environmental conditions (warm and cold) in the temperate and boreal ecozones of Quebec. Our goal was to disentangle the genetic from the environmental effects and understand their interaction for bud set, frost tolerance and non-structural carbohydrate content, and detect potential clinal variation in those traits.

Thus, this study should contribute crucial information to help minimize the risks of frost damage related to seed source transfer under intraspecific assisted population migration programs and support the development of new seed transfer models better able to cope with predicted climatic conditions, thus maintaining optimal productivity in future reforestation efforts in the context of climate change.

## MATERIAL AND METHODS

### Study area, seed sources and testing site conditions

In early 2010, seeds were collected in six first-generation white spruce seed orchards that were made up of regionally selected plus-trees (number of plus-trees varies between 187 and 284) and are among the most widely used in white spruce reforestation programs in Quebec (see Benomar et al., 2016, 2018 for more details). Seedlings from these six seed sources as well as from local seed sources (control) were planted between 2013 and 2015 at nine test sites distributed between latitudes 45°N and 49°N and longitudes 65°W and 79°W in the province of Quebec by the Ministère des ressources naturelles et des forêts du Québec (MRNF) (Otis Prud’homme et al., 2018).

These plantations were distributed along a mean annual temperature gradient of 5.8 °C. Two of these test sites with most contrasting mean annual temperatures were considered in this study, the southern Wendover site (planted in 2014) and the northern Rousseau site (established in 2015) (Figure 1). Each test site thus contained six transferred seed sources and the local seed source (Table 1). The photoperiod was estimated for each test site based on their latitude and longitude. The maximum difference in the daylight period between the northern and southern sites observed during the summer solstice was around half an hour (Supplementary Figure 1).

**Figure 1.**
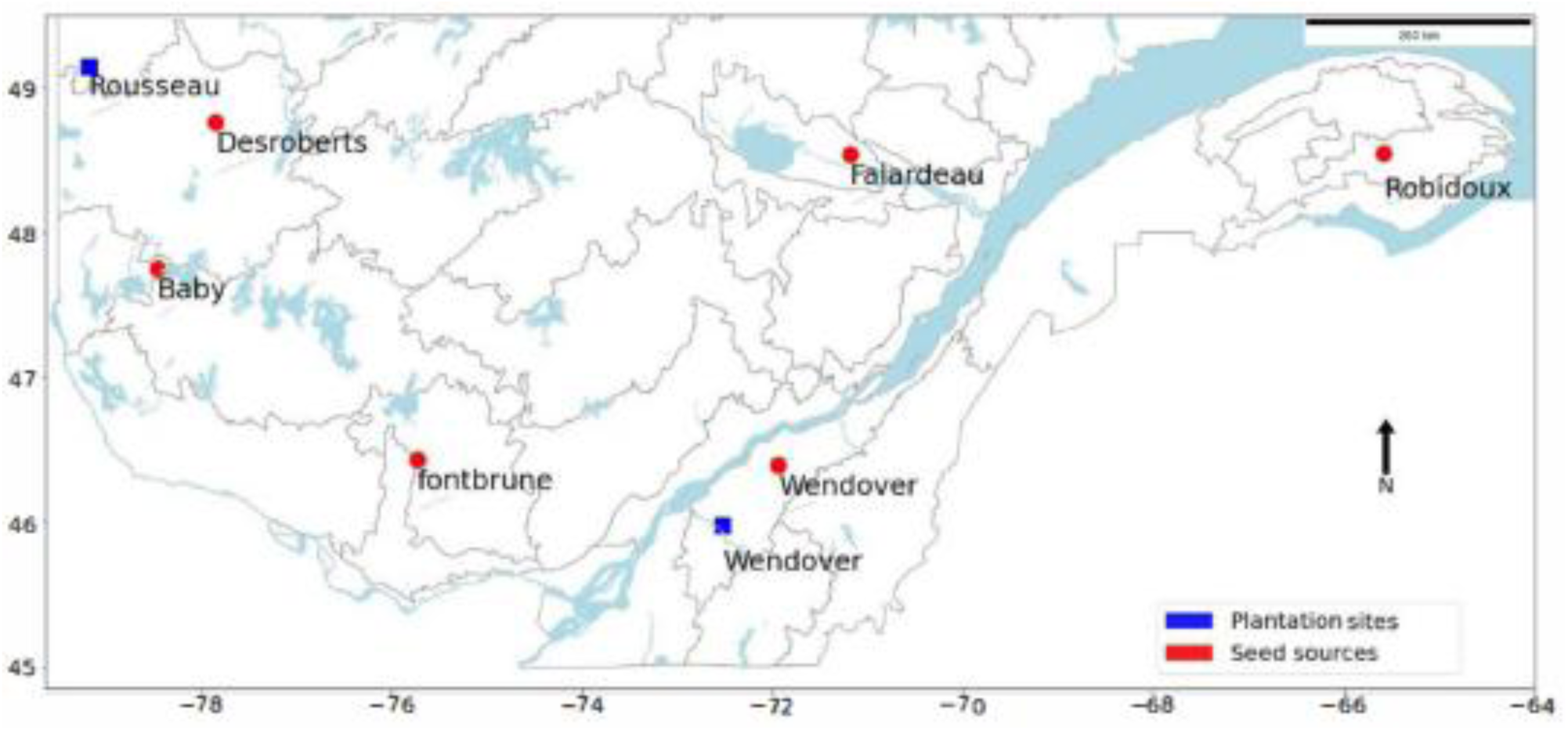
Geographical locations of seed sources and test sites.

**Table 1.**
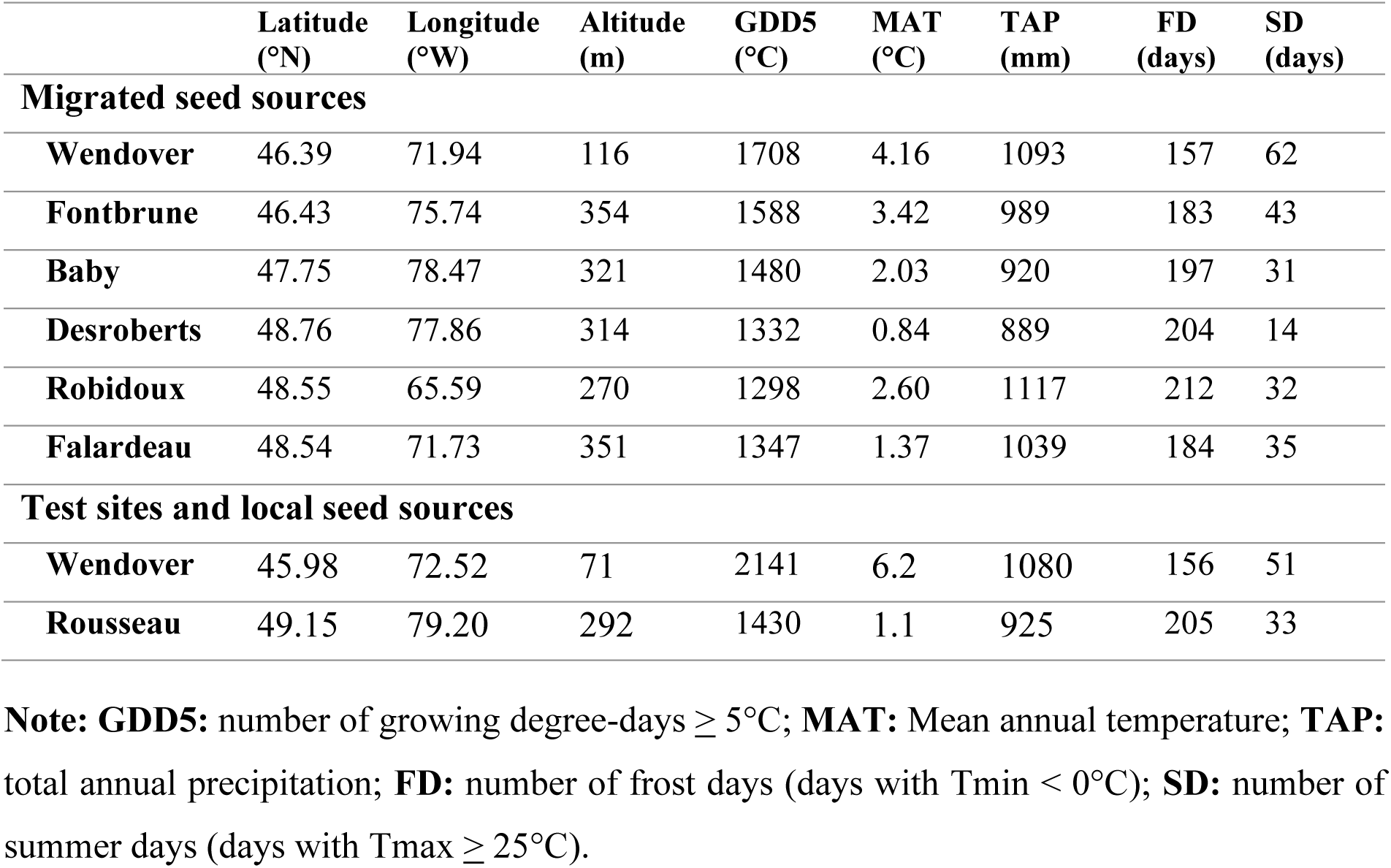
Geographical coordinates and bioclimatic normals of the origins of the six migrated white spruce seed sources (reference period 1940-1970) and of the two test sites (timeframe: 2010-2020). Climatic data was extracted using BIOSIM 10 (Régnière et al., 2014). The table was adapted from Benomar et al. (2016).

### Experimental design

Each plantation contained 4,032 white spruce trees organized in 4 randomized complete blocks (density of 2,000 trees/ha; spacing of 2.25 m x 2.25 m). Each block had 7 plots (each being occupied either by one of the 6 migrated seed sources or the local source), and each plot contained 144 trees (12 × 12). The outer rows were excluded from the measurements to avoid any plot border effect. Thus, only the central 64 trees per plot were considered for sampling and analysis.

### Climatic data

These seed orchards, referred to as seed sources herein, consisted of trees of regional origin (meaning they originated from natural populations located in the same bioclimatic domain in which the seed orchard was located). Considering a certain regional climatic homogeneity and the central location of this orchard, the climatic data origins used in this study correspond to those of the seed orchards. Seed source climatic data were obtained using the BIOSIM10 software (Régnière et al., 2014), which allows the extrapolation of temperature and precipitation from data collected in the most adjacent meteorological stations. The climate conditions of seed source origins were assessed using the local climate normals experienced by the parental generation from 1941 to 1970 as a seed source reference period, while the current climate (testing reference period) of each test site was assessed using the climatic average of test sites from 2010 to 2020. Multiple bioclimatic variables (https://www.climdex.org/learn/indices/) were derived from temperature and precipitation data using the climdex R package (Bronaugh, 2014) to assess the underlying mechanisms of adaptation and the interaction effects between seed sources and test sites.

### Bud set monitoring and frost damage

A follow-up of the different phenological stages of bud set was carried out on 20 trees/plot/block/site during the 2021 growing season. Observations were made weekly, with a one-week delay for the southern site given the phenological delay in the more temperate meridional region, from the end of June to early August. For each tree, the chronology of the average bud set stage made on several lateral shoots ranging from 7 to 10 shoots per tree, was recorded considering the different geographical exposures of branches and tree heights. The phenological stages were recorded using the white spruce phenology field guide (Dhont et al., 2010). The studied bud set stages are described as follows: Stage 0, no buds but elongation of the main stem may have already been completed; Stage 1, apical bud initiation with the presence of a white bud at the end of the main stem hidden under the needles; Stage 2, Beige bud with the scales that begin to cover the bud becoming beige and the size of the bud had increased; Stage 3, browning bud with the apical bud completely covered with browning scales and the volume of the apical bud had increased significantly; Stage 4, brown bud with the apical bud being completely brown and clearly visible and the needles of the whorl beginning to open outwards; Stage 5, open needles with the apical bud being clearly visible, with brown opaque and concave scales well-formed. To evaluate the damage from the late spring frost that occurred at the end of May 2021 in most of the Quebec territory (Benomar et al., 2022), the health state of trees was also recorded using a crown damage index, which was defined as the proportion of the crown of the trees affected by the late cold spell. Trees with more than 70% crown damage index were simply avoided.

### Evaluation of frost tolerance

A frequency analysis of the first fall frost was conducted over the last 30 years for each test site using the daily minimum temperature data. This analysis allowed us to identify the latest date by which the first fall frost typically occurs between the end of the bud set and the onset of frost. October 3^rd^, 2021 was identified as the best late sampling date to avoid any potential fall frost damage. For each seed source/block/site, composite samples of 24 shoots were randomly collected from six to eight trees. Shoots were stored in plastic bags previously labelled. Each bag contained a wet absorbent paper to provide a fresh environment to the samples and avoid their deterioration through direct contact with ice when placed in a cooler and sent to the MRNF laboratory in Quebec City for analysis the following day.

Frost tolerance was assessed by measuring electrolytic leakage from needles exposed to different freezing temperatures. The relative electrolytic conductivity (REC) is indicative of membrane leakage so the more the REC is important, the more important are the frost damages, with extensive cell membrane rupture. It is well known that low REC values reflect high frost tolerance (eg. Lamhamedi et al., 2005, 2022) and, ultimately, that seed sources are more likely to adapt and grow under these freezing temperatures. The frost tolerance test consisted of applying 6 temperature levels ranging from T_0_ = 4 °C (control) to T_5_ = -16 °C with a step of 4 °C using a programmable freezer (Model T20RS Tenney Environmental Inc., Williamsport, PA, USA). Each temperature level lasted 4 hours, and then the temperature was lowered at a rate of 1 °C·h^−1^; once the target temperature was attained, it was held constant for 4 hours, and the REC was estimated according to the protocol proposed by Lamhamedi et al. (2005, 2022). Briefly, for each site and each freezing temperature, four shoots were used for each seed source in each block. After having undergone the 5 levels of freezing temperature (T_1_ = 0 °C, T_2_ = -4 °C, T_3_ = -8 °C, T_4_ = -12 °C, and T_5_ = -16 °C) as samples of T_0_ were not subjected to freezing treatment, the samples were then immersed in 100 ml of demineralized water overnight at a temperature of 4 °C. After saturation, the release of electrolytes (EC1 measured in μSiemens/cm) in the solution of each sample was measured using a conductivity meter (model 160, Orion Research Inc., Boston, MA, USA). Afterwards, the maximum release of ions (EC2, µS/cm) was measured after placing the sample in the autoclave at 121 °C for 15 min and overnighting at 4 °C. Two frost tolerance indices were calculated as follow:

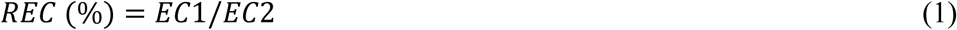

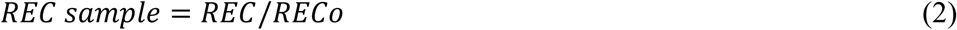

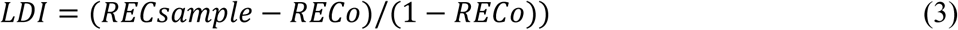

where EC1 is the measured electrolytic conductivity at a given freezing temperature at the beginning of the test; EC2 is the measured electrolytic conductivity at the end of the test for the same freezing temperature (1). REC sample is the relative conductivity at a given freezing temperature (2). RECo is the relative conductivity at reference temperature 4 °C; and LDI is the leaf damage index (3).

### Tree height measurements

To assess the growth performance of seed sources and detect any possible frost effects on their growth, total tree height at 7 years after plantation was measured in both sites, and mortality was recorded for all seed sources and within each test site. Due to the 1-year difference of trees’ age between sites, the height was adjusted for 6-year-old trees by adding the shoot length of 2021 to the total height of the same year to allow a comparison of height between test sites as a proxy for the same age.

### Non-structural carbohydrates (NSC) analyses

To examine variations in non-structural carbohydrate (NSC) content towards the end of the growing season in relation to cold hardiness and frost tolerance, the sampling of shoots was carried out on two dates during the fall of 2021: the first sampling day in early September and the second one in early October. For each seed source, three trees of different height classes (small, medium, and high) were sampled. Synchronizing NSC and sampling for frost tolerance assessment was designed to determine the relationship between frost tolerance and sugar metabolism. After collection, the samples were quickly stored in a cooler containing dry ice. Then, they were freeze-dried to preserve the integrity and quality of the sugars. Finally, they were sent to the TransBIOTech laboratory (Lévis, QC, Canada) to determine the NSC content using an ultra-performance liquid chromatography-evaporative light scattering detector (UPLC-ELSD). These analyses focused on the reportedly most important sugars for cold acclimation, that is, raffinose, glucose, sucrose, fructose, and pinitol (Lintunen et al., 2016).

### Statistical analyses

The odd ratio of annual frost probability during the growing season between test sites (Supplementary Figure 2) showed an extreme inter-annual variation of frost probability during the after-planting years in the northern test site. For estimation, the growing season was defined as between June 1^st^ and October 31^st^ of each year, and the number of frost days for each year was estimated as the number of days with daily minimum temperature < 0 °C during this period (Supplementary Figure 2).

To characterize the climate conditions of origins of seed sources and actual conditions at test sites, the annual odd ratio of frost probability during the growing season and the length of the growing season were calculated using the *climdex.gsl* function from the climdex R package (Bronaugh, 2014). These two variables were fitted separately to simple linear models to evaluate the differences between seed sources and test sites. The significantly different groups were mapped for both variables along the north-south geographic gradient.

Normality and homogeneity of variance assumptions were assessed and validated prior to the analysis of variance (ANOVA) for all fitted linear models in this study. These models were fitted using the *lm* and *aov* functions from the stats R package (R Core Team 2022). The presence of significant differences was followed by a *post hoc* analysis using Tukey’s Honest Significant Difference test from stats R package (R Core Team 2022) to identify the best seed sources for growth and frost tolerance. The R-square coefficient was used to estimate the explained percentage of the total variance using the Performance R package (Lüdecke et al., 2021).

The likelihood probability of bud set stage was modeled as a function of the day of the year, the test site and the seed source (model 4 below) by an ordinal logistic regression using the *polr* function of the MASS R package (Venables & Ripley, 2002). The results were visualized using the ggeffects R package (Lüdecke, 2018) to highlight the variation of bud set probability over time between seed sources and test sites. The model formula is as follows:

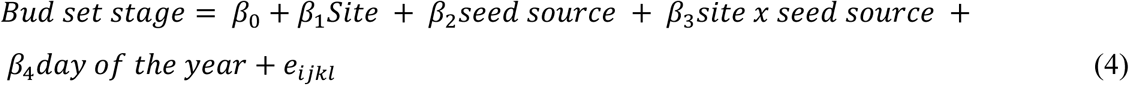

Successive stages correlations were examined as well (Supplementary Table S1).

The percentage of damaged terminal shoots and survival rate were fitted to a linear mixed model with fixed effects for seed sources, test sites and their interaction, and as a random effect for the blocks nested within site effect as follows:

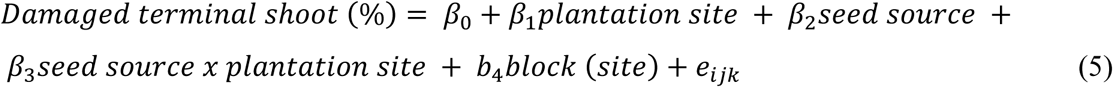

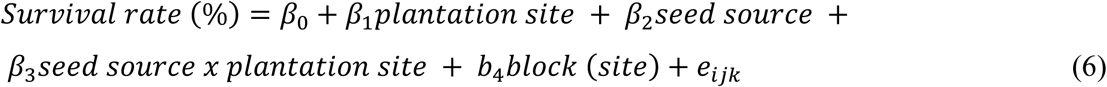

To evaluate seed sources’ frost tolerance, two linear models were fitted to REC and LDI vs. test site, seed source, freezing temperature, and all their interactions. The model formula is as follow for both frost tolerance indices:

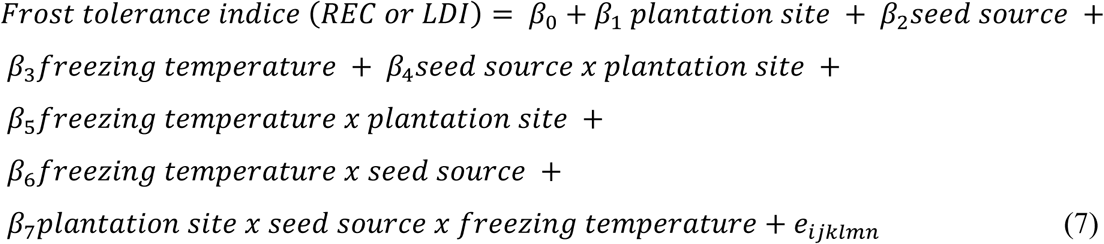

where *β*₀–*β*₇ are fixed-effect parameters to be estimated, and eᵢⱼₖₗₘₙ represents the residual error term. Total height was analyzed using the model in (5), while NSC content was analyzed using the model in (7) after replacing the freezing temperature by sampling date.

To determine the relationship between the climate of seed sources’ origins, the climate of test sites, frost tolerance indices, NSC content, tree height and bioclimatic variables, a correlation analysis was conducted on the average values of seed sources alone, then with the average values of test sites for all 24 variables. Similar correlations were grouped together using the Ward complete hierarchical clustering method (Ward, 1963). The resulting clusters were visualized in two heatmaps (with and without test sites’ average values) with significance levels using corrplot R package (Taiyun & Viliam, 2021). All significant correlations of frost tolerance indices were scatter-plotted to highlight linear relationships of seed sources’ origins and the test sites for bioclimate, frost tolerance, height and NSC content. To determine the discriminating factors of the studied seed sources and test sites, bioclimatic variables were scaled and centered, and then a principal component analysis (PCA) was fitted using the NIPALS (Nonlinear Iterative Partial Least Squares) method using the pcamethods R package (Stacklies et al., 2007). The Factoextra R package (Kassambara & Mundt, 2020) was used for visualization.

## RESULTS

### Climate characterization of seed source origins and test sites Length of the growing season and frost probability at seed source origins and test sites

The average length of the growing season, calculated over the reference period (1940-1970), showed that the Wendover seed source had the longest growing season, around 150 days (Figure 2). The northwestern seed sources Baby and Desroberts exhibited the shortest growing seasons of around 100 days. The Wendover seed source was significantly different from these two northwestern seed sources. The northeastern Robidoux, northern Falardeau and southwestern Fontbrune seed sources exhibited a medium average length of the growing season, ranging from 115 to 130 days. These three seed sources were statistically similar and were clustered into an intermediate group between the southern and the northwestern seed sources for the length of the growing season (Figure 2). According to a different reference period (testing reference period) as compared to seed sources, the test sites showed significant differences in their average length of growing season (for the contemporary 2010-2020 period), with 165 days for the northern site Rousseau, and 200 days for the southern site Wendover (Figure 2), a gain of 50 days in the growing season at Wendover between the two periods considered, indicating the impact of global warming underway in the temperate ecozone of southern Quebec.

**Figure 2.**
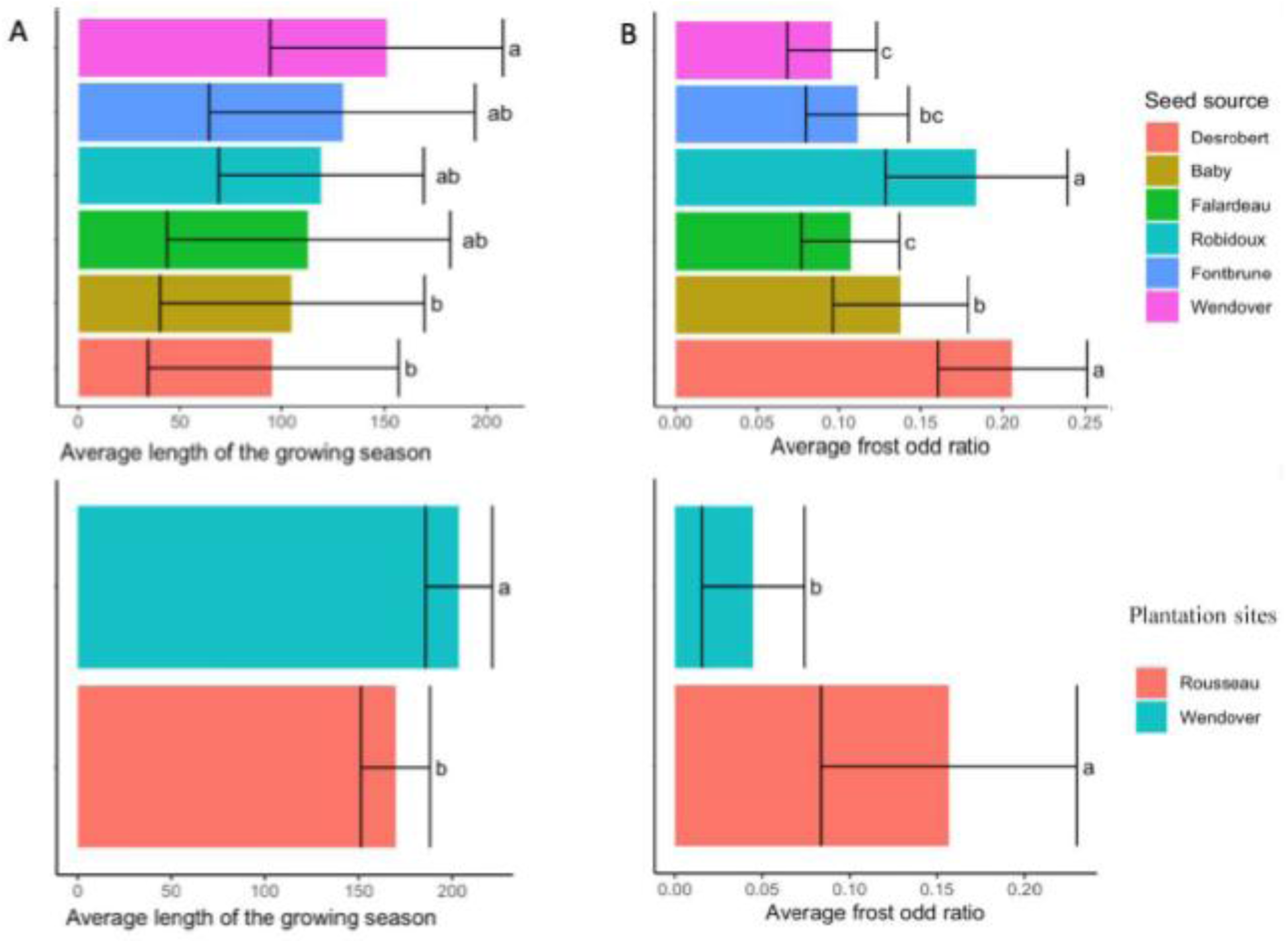
Means and standard deviations of the length of growing season (A) and frost odd ratio probability (B) for seed source origins and test sites, with actual climate data (2010-2020, as reference period) for test sites and data of climates of origins (1940-1970) for seed sources. The beginning of the length of the growing season was determined as 6 consecutive days with temperature ≥ 5 °C, from June 1^st^ to October 31^st^. Seed sources and test sites are listed from north to south.

Regarding the average frost odd ratio probability during the growing season, the Wendover, Fontbrune, and Falardeau seed sources showed the lowest probabilities, with values around 0.1 (Figure 2). The probability was significantly higher for the Baby seed source at mid-latitude, with an average of 0.13. The highest probabilities were observed for Robidoux (northeast) and Desroberts (northwest) seed sources, ranging from 0.18 to 0.21. The Fontbrune southern seed source belonged to an intermediate group that combines the c and b Tukey groups for the frost odd ratio probability (Figure 2), which are also represented by the southern Wendover and northern Baby seed sources. Regarding the test sites, significant differences were recorded, with 0.05 average frost probability for the southern site, and 0.16 for the northern site (Figure 2).

When considering the length of the growing season of seed sources and the average frost odd ratio probability together, an expected geographic association was found. The frost odd ratio probability increased from south to north and, as expected, the opposite trend was observed for the length of growing season (Figure 2).

### Chronology of bud set phenology

The chronology of bud set of the studied seed sources followed a sigmoidal curve starting from 0 (last week of June) to 5 (final bud set stage: 1^st^ week of August). The trees in the southern Wendover test site showed an earlier bud set for all stages and all seed sources compared to those planted in the north, and all seed sources reached the final bud set stage at the same time in both sites (Figure 3a). However, the last stage of bud set was similar for all seed sources in each site. Thus, the bud set period of the studied seed sources was longer at the southern test site. On the other hand, the average normal GDD5 to initiate bud set was higher for the Wendover southern test site, around 700 growing degree-days, versus 490 growing degree-days for the Rousseau northern test site (Figure 3b).

**Figure 3.**
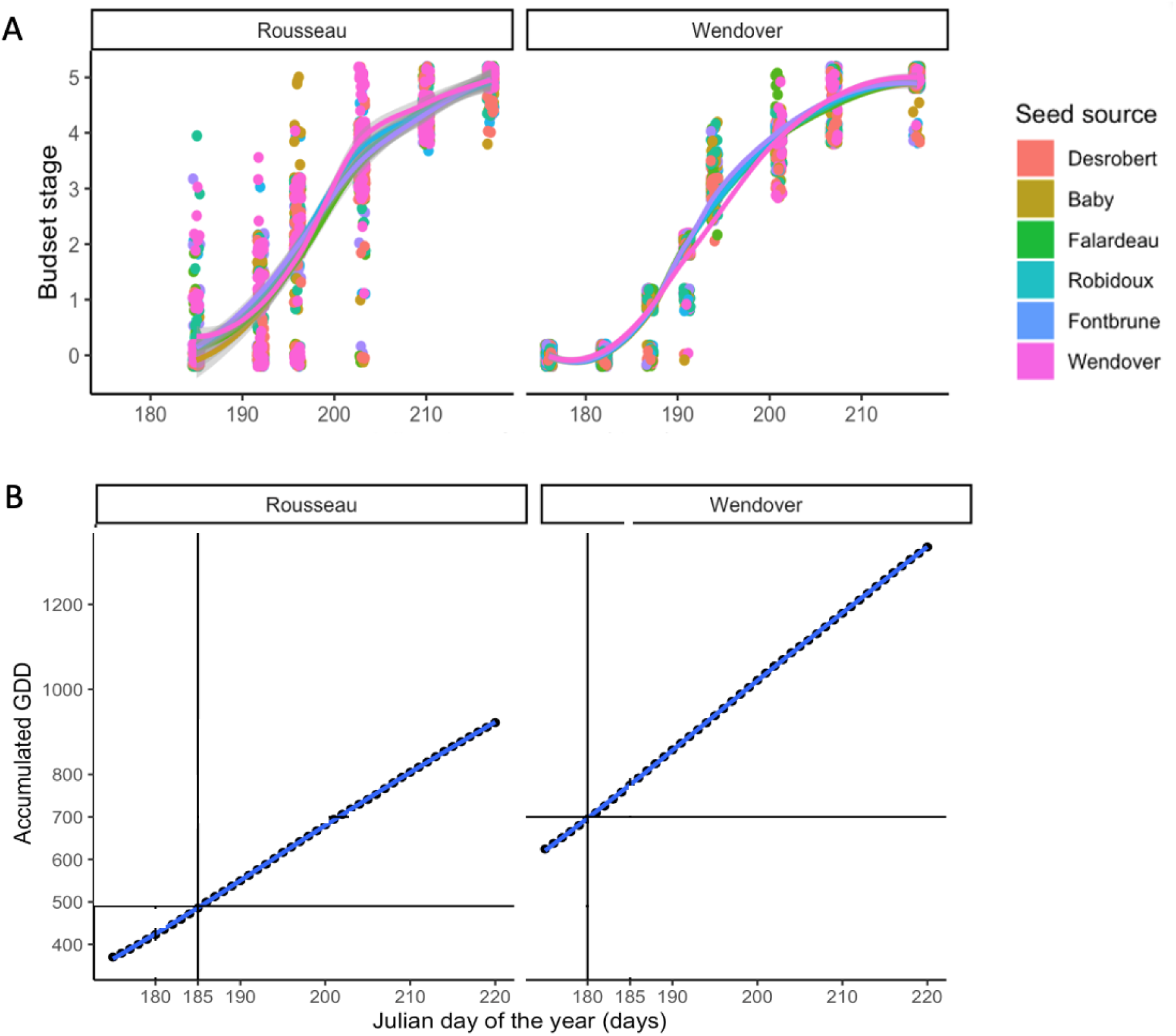
The chronology of bud set of the studied seed sources for each of the northern Rousseau and southern Wendover test sites. (A) presented with the corresponding average normal GDD (cumulative growing degree-days ≥ 5 °C calculated over a 10-year period (2010-2020)). (B) Seed sources are listed from south to north. Note: GDD5=0, if (T_min_ + T_max_) < 5°C; and GDD5 = (T_min_+T_max_)/2 -5, if T_min_ + T_max_ ≥ 5°C.

### Modeling bud set phenology of seed sources at test sites

The ordinal logistic regression model of bud set explained 63% of the total variation according to McFadden *R^2^* coefficient (Supplementary Table S1). Bud set initiation was earlier in the southern test site and for all bud set stages. It highlights a highly significant effect of the day of the year on the bud set transition (Figure 4A) and less interference were recorded for stages 0, 4, and 5. However, stages 1, 2, and 3 showed moderate occurrence probability (with a maximum of 0.5) and higher interference between stages in the time frame between 185 and 205 Julian days of the year (Figure 4A). Regarding variation among seed sources, the same pattern was recorded for stages 0, 4, and 5, with higher occurrence probability at the beginning and the end of the bud set cycle. Higher variation was recorded for stages 1, 2, 3, and 4 during the time frame between 190 and 205 Julian days of the year, with moderate probabilities and higher interference between bud set stages (Figure 4B). All seed sources’ coefficients were significantly different to those of the Wendover seed source except for Fontbrune, another southern seed source (Supplementary Table S1). A highly significant G×E interaction implicated the southern Wendover test site and the seed sources Desroberts, Baby, and the local seed source of that test site (Supplementary Table S1).

**Figure 4.**
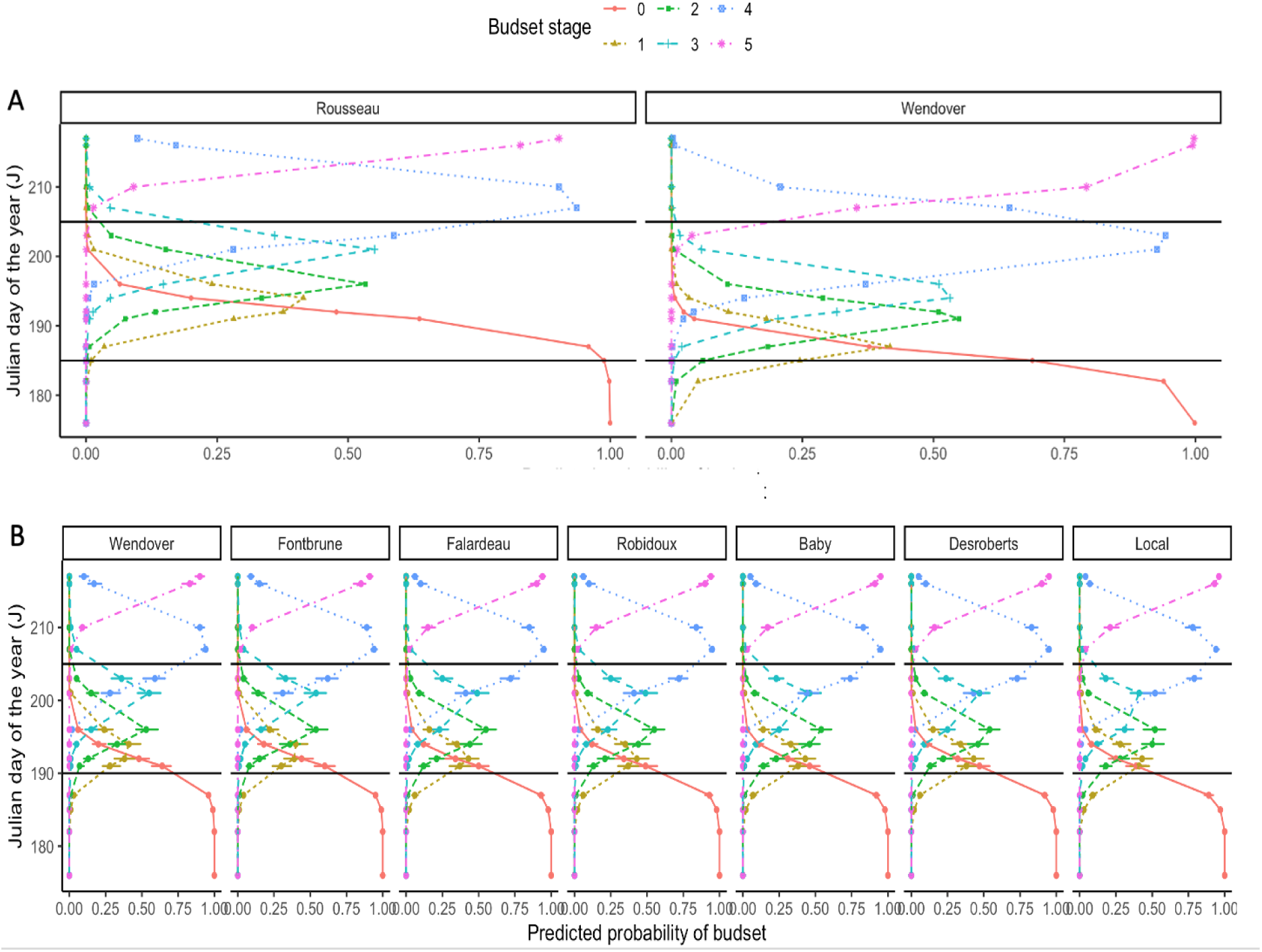
Predicted probability distribution and confidence intervals of bud set stages of white spruce by test site and seed source through time: A) northern Rousseau and southern Wendover test sites, and B) the seven seed sources, including the local seed source for each test sites.

### Levels of frost damage of terminal shoots and survival rate of seed sources at test sites

The percentage of trees of each seed source showing shoot frost damage in both test sites following the late cold spell of 2021 (Benomar et al., 2022) is presented in Figure 5. Most seed sources planted in the southern Wendover test site showed moderate to high damages to terminal shoots, with values ranging from 50% to 80% (particularly for the local seed source). This site also showed high variability compared to the northern Rousseau test site where most seed sources had a percentage of damaged terminal shoots of around 50%. The lowest level of shoot damage was recorded for the Baby seed source (35%) in the northern test site. Little significant differences were recorded among seed sources and between test sites for shoot frost damage and survival rate (Figure 5).

**Figure 5.**
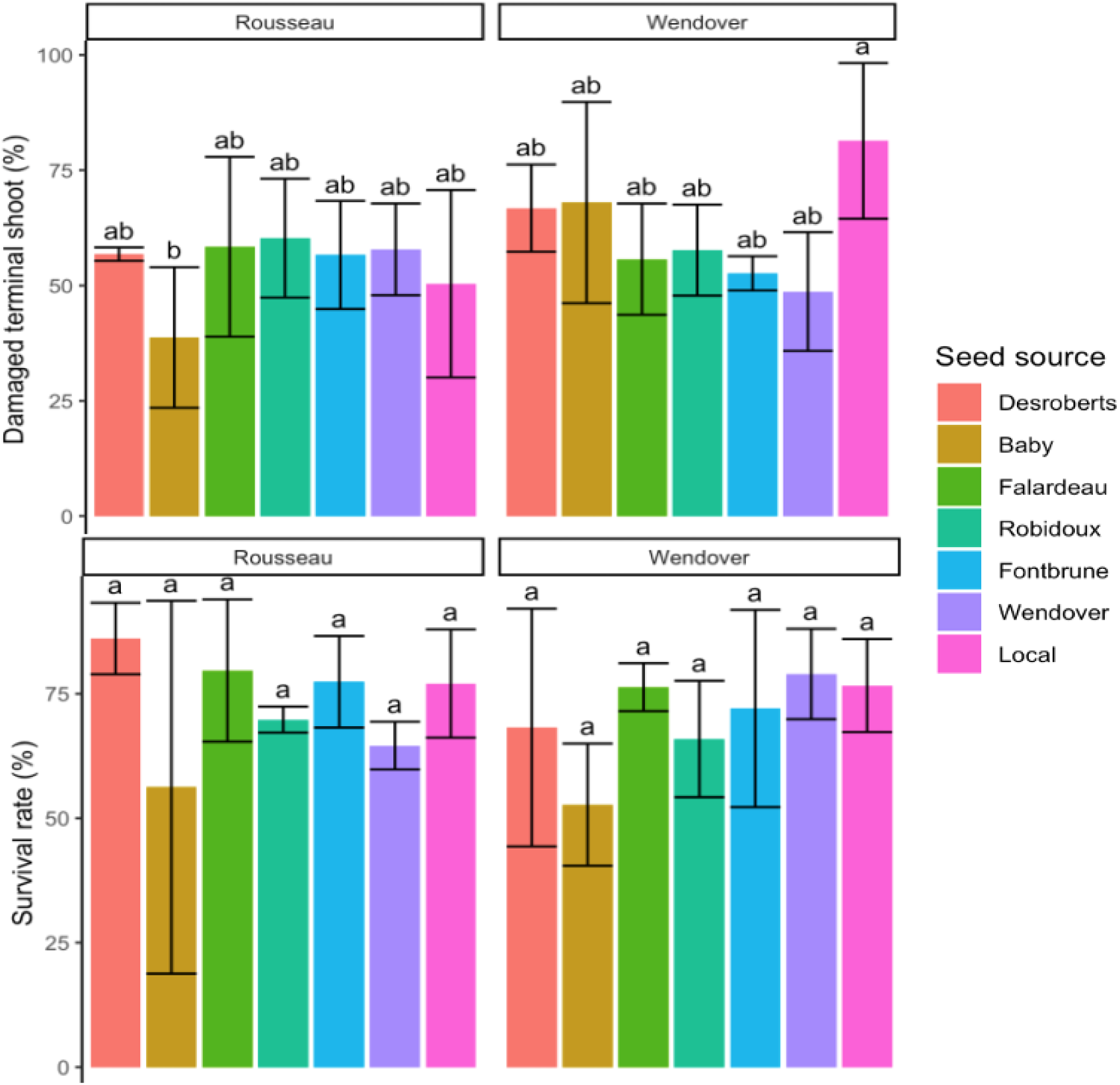
Distribution of the average percentage of damaged terminal shoots and percentage of live trees for all seed sources within each test site.

### Analysis of variance of frost tolerance, height, and detected NSC compounds

Relative electrolytic conductivity (REC) and leaf damage index (LDI) were significantly influenced by the test site, the freezing temperature, and their interaction between both factors (Table 2). However, only REC showed an additional significant effect of the triple interaction between test site, seed source, and freezing temperature. For tree height, the seed source, test site, and their interaction effects were highly significant. For NSC compounds, the test site effect was significant for all analyzed sugars. Seed source and sampling date effects were only significant for sucrose content. However, the interaction between test site and sampling date was significant for fructose, sucrose, and glucose contents (Table 2).

**Table 2.**
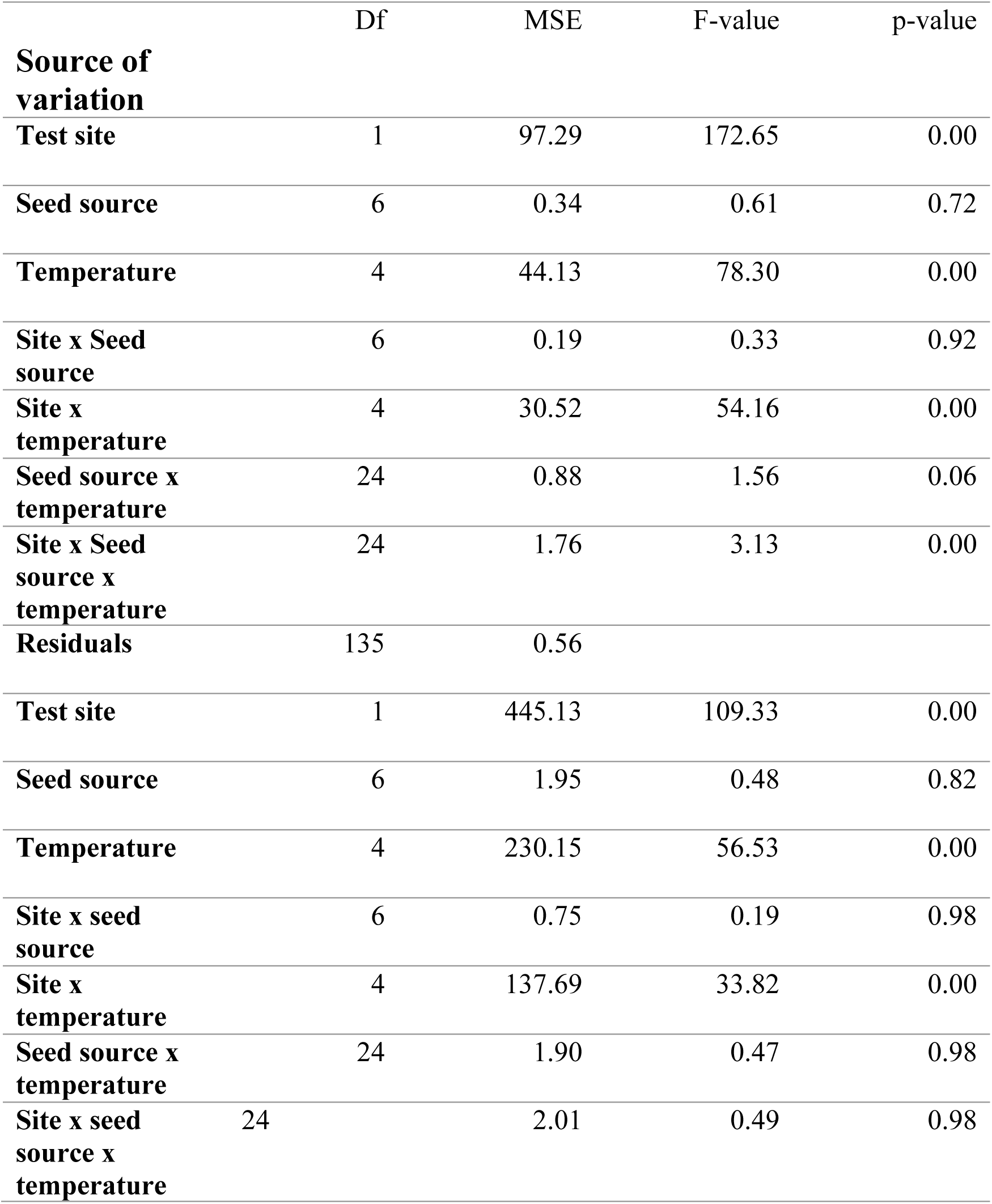
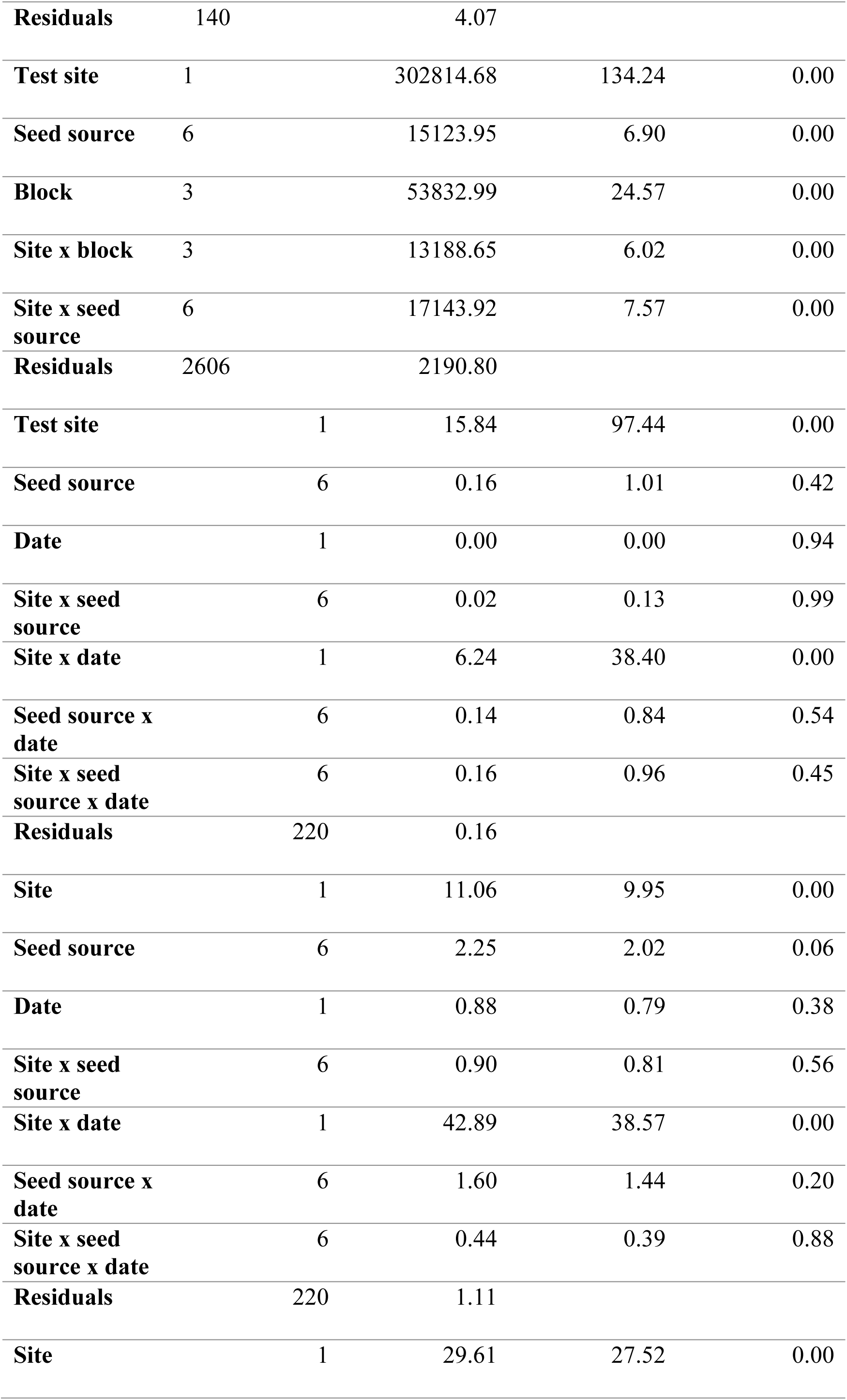
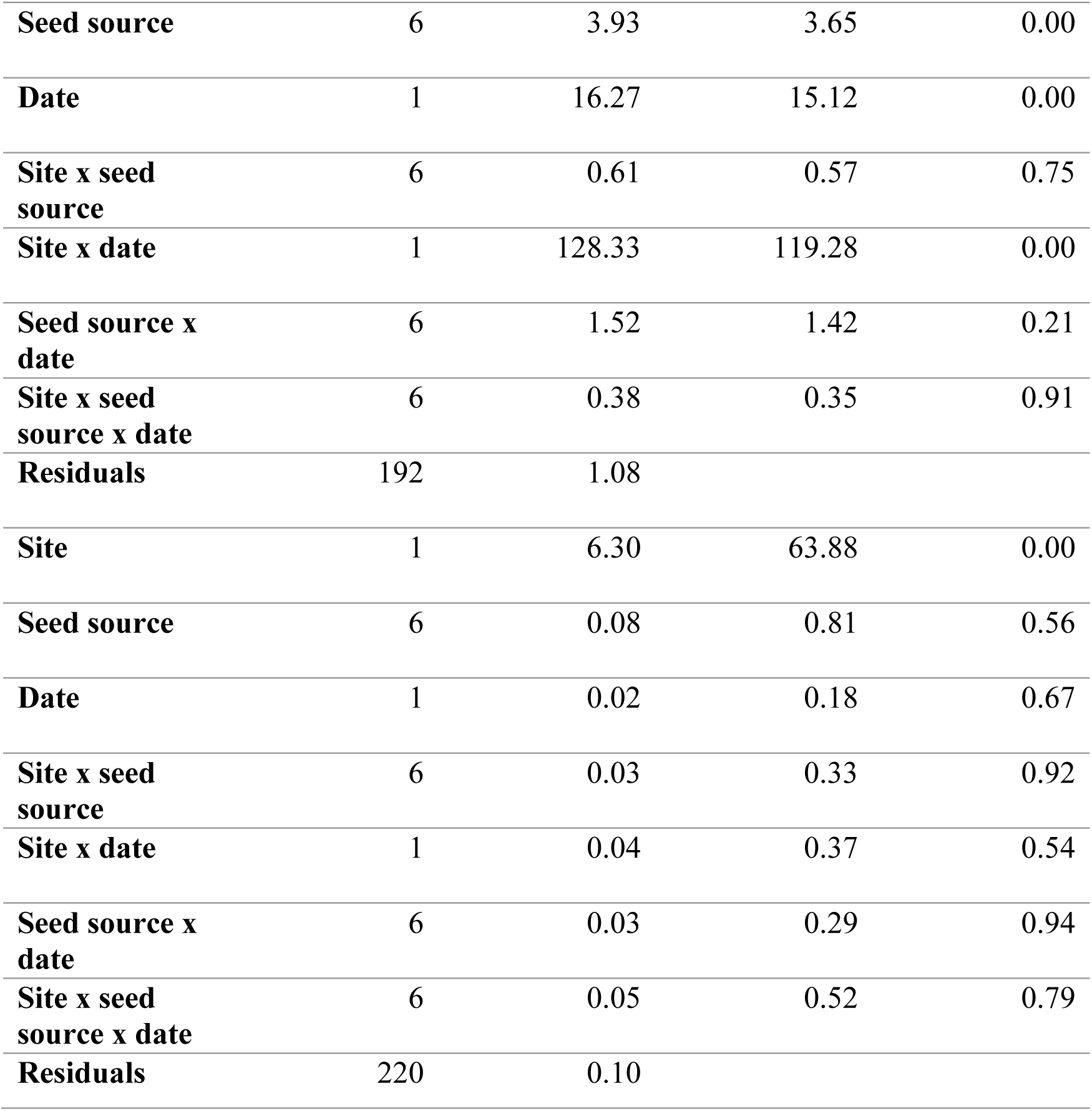
Summary statistics of the analysis of variance (ANOVA) of frost tolerance indices, tree’s height, and NSC content. A: ANOVA for frost tolerance (REC, relative electrolyte conductivity) considering the effects of test site, seed source, freezing temperature, and their possible interactions. B: ANOVA for leaf damage index (LDI) considering the effects of test site, seed source, freezing temperature, and their possible interactions C: ANOVA for 7-year height considering the effects of test site, seed source, and their interaction. D to G: ANOVAs for NSC sugar contents considering the effects of test site, seed source, sampling date, and their interaction (D, fructose; E, glucose; F, sucrose; G, pinnitol). Abbreviations used in the table are described as follows: Df, degrees of freedom; MSE, Mean-square error; F-value, Fisher’s test value; p-value, probability level of significance.

### Frost tolerance indices: REC and LDI of seed sources at test sites

Significant differences in REC and LDI were detected by ANOVA and more details on the trends observed are presented here. The REC boxplots of the studied seed sources showed an increasing pattern with high values (low frost tolerance) under -12 °C and -16 °C in the southern Wendover test site with Falardeau and Wendover seed sources standing out from the others for freezing temperatures of -16 °C (Figure 6A). In contrast, most of the seed sources in the northern Rousseau test site showed a more uniform pattern following the decrease of freezing temperature. Thus, all seed sources planted in the northern site showed low electrolytic conductivity, except for Falardeau, which reflects their capacity to tolerate higher freezing temperatures.

**Figure 6.**
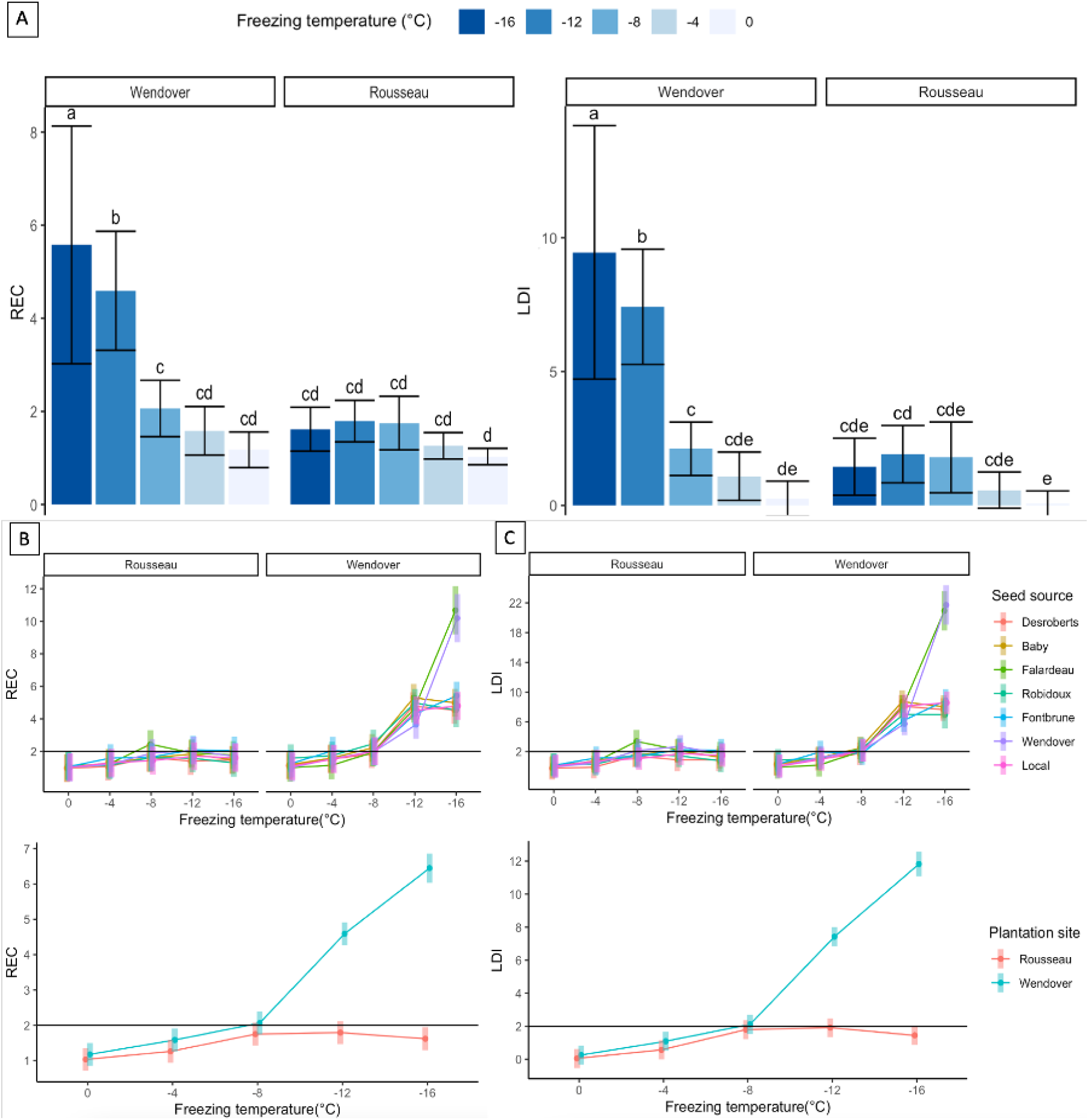
Estimated means and standard deviations with confidence intervals of relative electrolytic conductivity (REC) and leaf damage index (LDI) of the studied seed sources within each test site (Rousseau and Wendover) sampled October 3^rd^, 2021 and for the five tested freezing temperatures (A), (B) and (C) represent predicted REC and LDI under the effects of seed source, test site, freezing temperature, and their interactions. Seed sources are listed from north to south.

Similar patterns were recorded for LDI, highlighting its strong correlation with REC. Seed sources with LDI values around 2 (black horizontal line, Figure 6C) showed no apparent frost damage, which was especially so for the Desroberts northern seed source and the Robidoux eastern seed source (Figure 6B, & C). The observed patterns for REC and LDI were statistically significant (Table 2A, & B), as confirmed by the Tukey test (Figure 6), highlighting three different groups, mainly for the southern test site (Figure 6A) based on freezing temperature: - 16 °C, -12 °C, -8 °C, and 0 °C. These groups also allowed identifying the most vulnerable seed sources (Falardeau and Wendover) under low-freezing temperatures (Figure 6B, & C).

### Height variation within and among seed sources at test sites

Given that significant effects were detected by ANOVA for 7-year tree height, we looked at the variation in more details. The distribution of 7-year tree height showed higher variation among seed sources at the Wendover southern test site compared to the Rousseau northern plantation site (Figure 7A). The average seed source height in the northern plantation site ranged from 120 cm to 145 cm while it ranged from 130 cm to 185 cm in the southern plantation site. The northeastern Robidoux seed source harboured the lowest average tree height in the southern test site, while the Fontbrune southern seed source had the highest average tree height in the same site (Figure 7B). The latter was significantly higher than the other studied seed sources planted in the south (Table 2). However, the Desroberts northern and Wendover southern seed sources showed almost similar average tree height. At the northern test site, no significant differences were recorded between seed sources for 7-year tree height (Figure 7A, & B).

**Figure 7.**
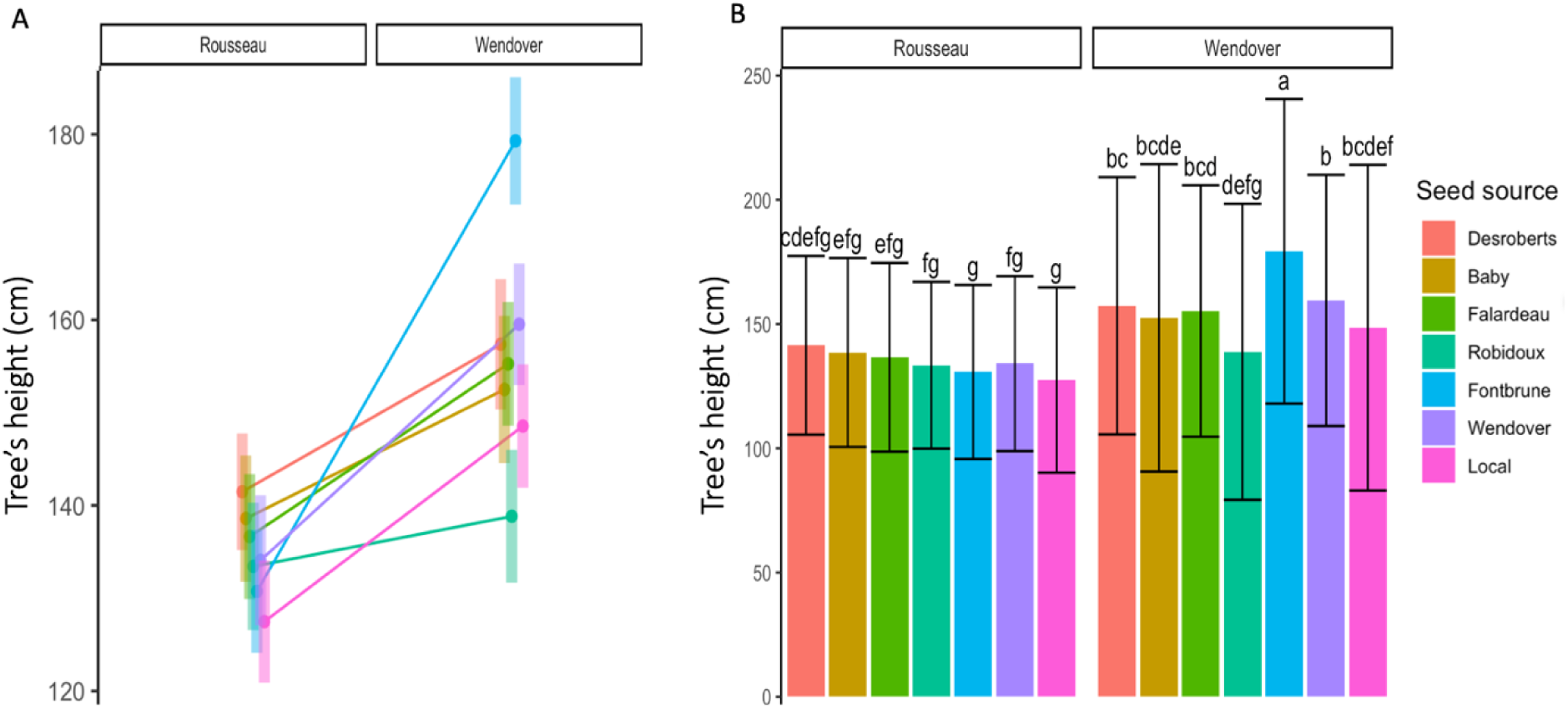
Distribution of means, standard deviations, and confidence intervals of 7-year tree height (cm) for all studied seed sources in the northern Rousseau and southern Wendover test sites. A) Represents the confidence intervals of predicted 7-year height from (Table 2) and B) represents the Tukey test results with statistical differences and grouping letters; Seed sources are listed from north to south.

Little interaction with plantation sites was observed for the Wendover, Falardeau, and Baby seed sources. In contrast, the southern Fontbrune seed source showed an important increase in tree height, from 130 cm at the northern Rousseau site to 180 cm at the southern Wendover site, highlighting its positive response to the warmer conditions in the south.

### Dynamics of non-structural carbohydrates (NSC)

Given that significant effects were detected by ANOVA for NSC, we further investigated their variation in more details. The chemical analysis of NSC compounds was conducted for four major sugars: fructose, glucose, sucrose, and pinitol. Following the variation in NSC content, many spatio-temporal patterns were recorded for fructose, glucose, and sucrose content (Figure 8A, B, C, & E) at both test sites and for both sampling dates. From September to October, fructose and glucose contents increased in the southern and decreased in the northern test site (Figure 8A, B, & E), while an opposite pattern was observed for sucrose content (Figure 8C & E). However, pinitol content showed significant differences only between test sites, with higher values in the southern test site (Figure 8D). A significant seed source effect was observed only for sucrose content with Robidoux showing the lowest levels, while Desroberts, Baby, and Wendover showed the highest ones.

**Figure 8.**
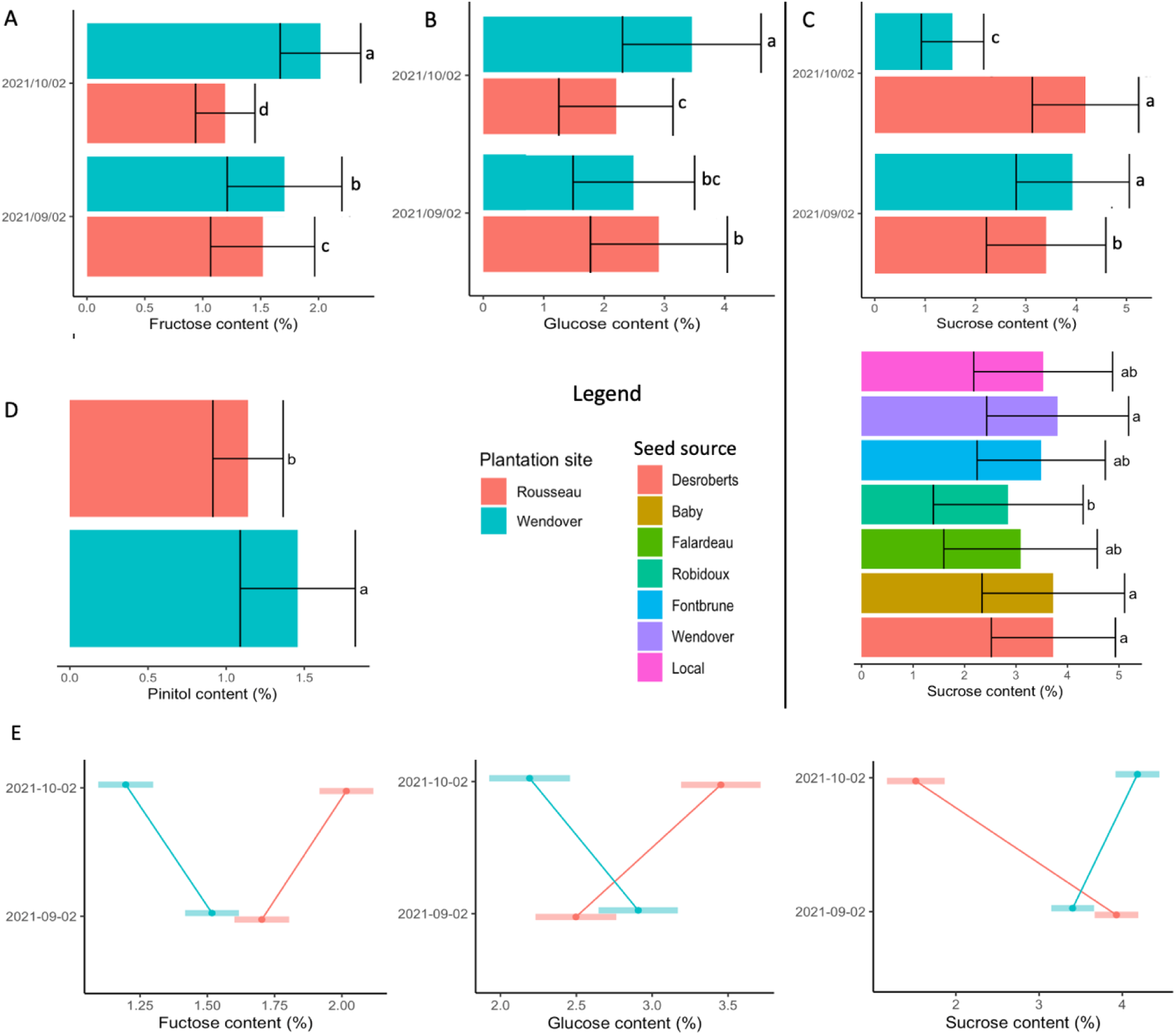
Means, standard deviations, and confidence intervals of interactions for all significant differences of non-structural carbohydrates (NSC) content detected between test site, seed source, sampling date, and their interactions. A) Fructose content, B) Glucose content, C) Sucrose content, D) Pinitol content, E) Spatio-temporal interactions of fructose, glucose and sucrose content. Seed sources and test sites are listed from north to south. The y axis of subplots colored by plantation site represents the sampling date.

### Correlation heatmaps of non-structural carbohydrates, height, frost tolerance indices, and bioclimatic variables at test sites and seed source origins

The correlation analysis of the mean values of all variables, taken pairwise, showed multiple associations between functional traits, growth, climates of seed source origins, and actual climates at test sites (Figure 9A). The hierarchical clustering of similar correlations resulted in six distinct groups, among which four were highly significant clusters (Pearson test, *P <* 0.05). The first group was characterized by variables such as the number of frost and icing days, the maximum length of the wet spell, and the daily temperature range describing association related to thermal variation and humidity conditions. The second significant group included pinitol and fructose contents, frost tolerance indices, and tree height. It described associations between ecophysiological traits and growth. The third group was more related to climatic conditions during the growing season, characterized mainly by maximal extremes of daily temperature and the number of summer days, which differentiate well the northern from the southern test site and the climates of seed sources origins. The fourth group was characterized mainly by precipitation variables and the minimum values of daily maximum and minimum temperatures, highlighting an association between the thermal amplitude and precipitation. When excluding the averages of the test sites, many biologically significant correlations were detected, highlighting the important role of ecophysiological plasticity in the clustering patterns of associations between variables (Figure 9B).

**Figure 9.**
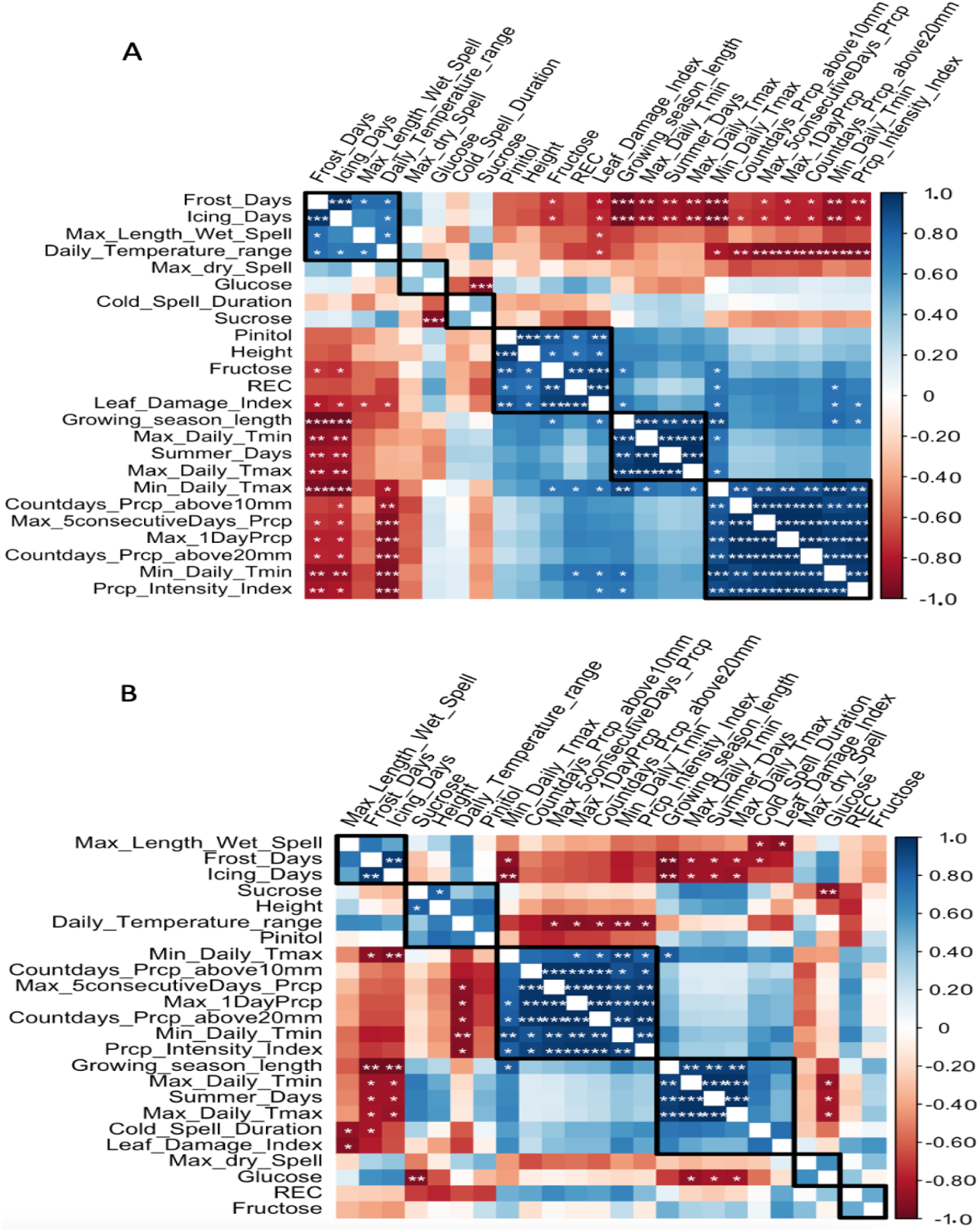
Pearson correlation heatmap of the averages of frost tolerance indices, non-structural carbohydrates content, tree height, bioclimatic variables of seed source origins and test sites (A) and the same correlation heatmap without the test site averages (B). * *P* < 0.05, ** *P* < 0.01, *** *P* < 0.001. The squares were delineated using hierarchical clustering.

Many significant pairwise correlations were detected between frost tolerance indices, NSC content, tree height, and the bioclimatic variables of seed source origins (Figure 10). Hence, following the climatic gradient of seed source origins from north to south and from west to east, the average leaf damage index was negatively correlated with tree height. Shorter trees appeared to be more prone to frost damage than taller ones, which is likely due to cold air masses being physically closer to the ground during late cold spells (Benomar et al., 2022).

**Figure 10.**
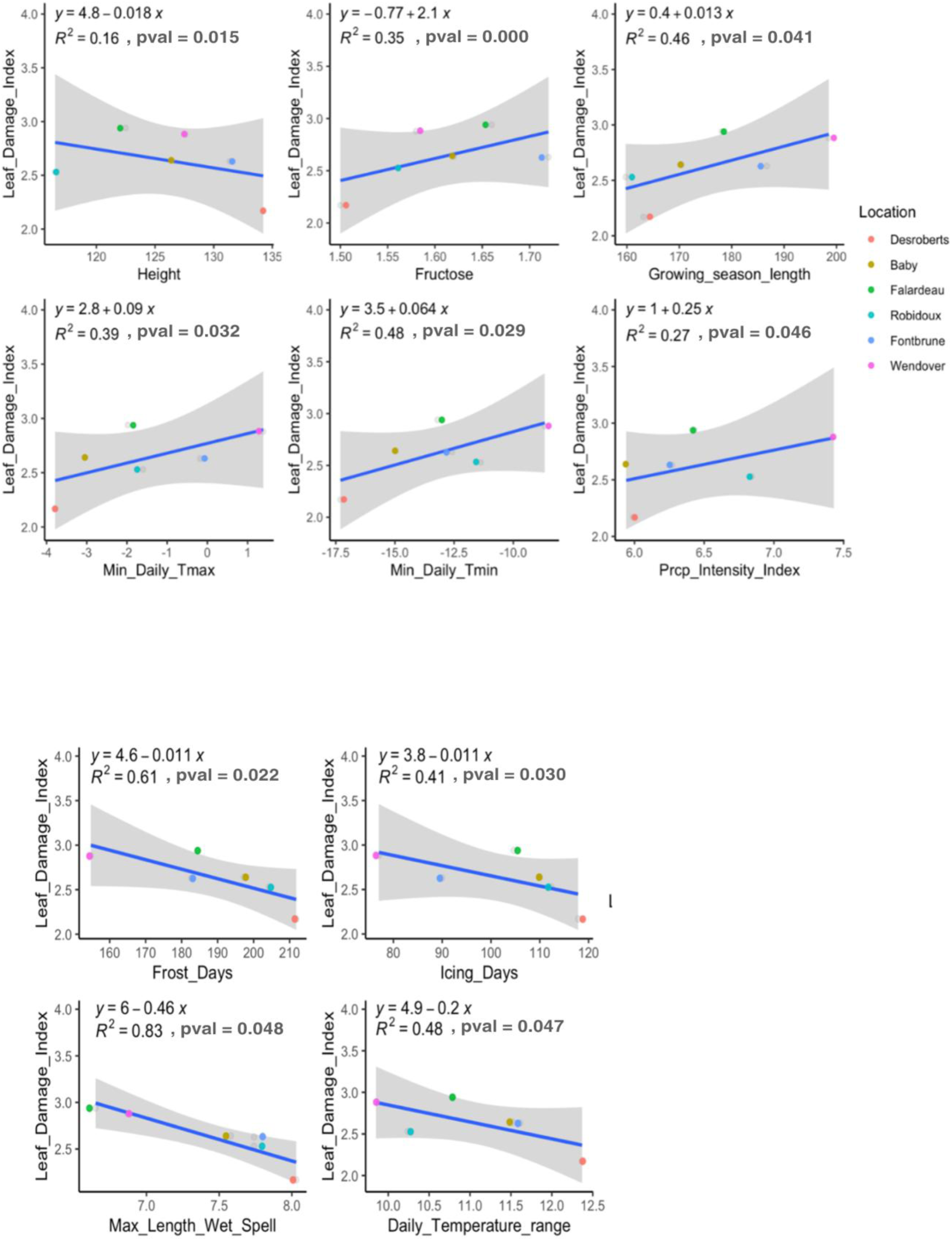
Relationship between seed sources average leaf damage index and non-structural carbohydrates content, tree height, and bioclimatic variables of seed source origins.

Fructose content was positively correlated with the leaf damage index, with the lowest value being recorded for the northern seed source. The mid-latitude Falardeau and southern Wendover seed sources had a similar leaf damage index despite having slightly different fructose contents. The average fructose content explained 35% (*R^2^*) of the total variance in the leaf damage index (Figure 10).

Regarding the bioclimate variables, the average length of the growing season, the minimum values of daily minimum and maximum temperature, and the precipitation intensity index showed positive relationships with the average leaf damage index. They explained 46%, 39%, 48%, and 27% of the total variance in the leaf damage index, respectively.

The longer length of growing season was associated with higher leaf damage index (Figure 10), likely indicating a propensity for more southern seed sources to have suffered more damage for the 2021 late cold spell. The average leaf damage index was also positively correlated to the number of frost days and icing days, the maximum length of wet spell, and the daily temperature range value. They explained, respectively, 61%, 41%, 83%, and 48% of leaf damage index total variance. All observed relationships of leaf damage index to seed source origins climate generally followed the same north-south geographic gradient (Figure 10). An exception was for the Falardeau seed source, which appeared to be the most vulnerable and being out of the confidence interval of most of the fitted models (Figure 10). The high percent of explained total variance of our models indicates the differential local adaptation of the studied seed sources. The rest of the unexplained variance was attributed mainly to test sites and the test site x freezing temperature interaction, as identified previously for the leaf damage index (Figure 10).

Four significant correlations were observed for the average REC in relation to NSC content (fructose and pinitol), height, and leaf damage index (Supplementary Figure 4). An increasing pattern was observed for fructose content and leaf damage index, explaining respectively 25% and 27% of REC total variance (Supplementary Figure 4). Knowing that high values of REC reflect low frost tolerance, the Desroberts seed source exhibited the lowest REC value among seed sources, that is, the highest frost tolerance. Regarding the leaf damage index, the Desroberts seed source had the highest frost tolerance and lowest frost damage, while the Robidoux seed source showed the lowest frost tolerance with medium frost damage. The Wendover and Falardeau seed sources had the same level of leaf damage with low frost tolerance among seed sources. Height and pinitol content had a decreasing pattern in relation to REC and explaining, respectively, 59% and 49% of REC total variance (Supplementary Figure 4). The lowest REC value, thus highest frost tolerance, was observed for the most northern Desroberts seed source, which also showed the highest average tree height value; while the mid-latitude eastern Robidoux seed source had the highest REC value (indicating lowest frost tolerance) with smallest average tree height among seed sources (Supplementary Figure 4).

## DISCUSSION

Given the predicted climate change in temperate and boreal forests, some species will be exposed to environmental stresses that are likely to cause significant damage to local populations and trees with low plasticity and maladapted genetic background, especially with the increased frequency of extreme climate events under mid-northern latitudes (Benomar et al., 2022). Assisted population migration was suggested early on as a proactive approach for climate change mitigation (Andalo et al., 2005; Beaulieu & Rainville, 2005). It consists of relocating species within their actual natural range to environments experiencing climatic conditions similar to those to which they were locally adapted previous to climate change. In this study, the evaluation of six regional white spruce seed sources on two test sites located at the extremes of a climatic gradient from southern Wendover to northern Rousseau test sites made it possible to simulate the effect of climatic transfers in the temperate and boreal ecozones. Through the study of past and actual climate conditions, bud set phenology, growth, frost tolerance, and non-structural carbohydrate metabolism towards the end of the growing season, it was possible to gain knowledge on the genetic adaptation and plasticity of the studied seed sources in terms of growth and physiological responses to environmental changes.

### Length of the growing season and frost probability at the location of the seed sources and test sites

Going from the southern to northern locations for test sites and seed source origins, the analysis of variance showed a decreasing pattern of the growing season length while the frost probability recorded a significant increase. The two northernmost seed sources (eastern Robidoux and western Desroberts) showed higher frost probability (Figure 2B). This association was also confirmed by the principal component analysis of the climate of seed source origins and the actual climate of the test site (Supplementary Figure 3). Following the geographic gradient from south to north, principal component analysis showed that northern origins and test site location were primarily correlated with the number of frost and icing days, while the southern origins and test site location were predominantly correlated with the growing season length, summer days, and precipitation indices. Our findings regarding the association between the length of the growing season and frost occurrence events are consistent with those reported by Marquis et al. (2022). Additionally, the maps produced by Ouranos showing many frosts and growing season indices under different climate scenarios in Quebec (available on their climate platform: https://www.ouranos.ca/en/) also support our findings.

### Bud set phenology of seed sources in contrasted environments

Modeling the probability of bud set phenology through time allowed to identify the significant effects of the test sites, the seed sources, their interactions, and the day of the year effect (Supplementary Table S1). The detected highly significant effect of the test sites highlights the phenotypic plasticity in bud set expression. These results are aligned with those of Jill et al. (2016), highlighting the earlier initiation of bud set in more meridional locations. However, other studies have reported earlier bud set initiation in more northern locations (Li et al., 1993, 1997; Beaulieu et al., 2004; Lesser et al., 2004). These differences might be related to differences in seed source variation, as well as in local photoperiod and temperature, which represent the essential factors to initiate bud development during the fall and the subsequent dormancy (Delpierre et al. 2016).

Using the Wendover seed source as reference in the ordinal logistic regression, it was possible to show that most seed sources had highly significant effects compared to the Wendover seed source. There was only one exception, that is the Fontbrune seed source, reflecting the adaptive capacity of the studied seed sources. Our findings are consistent with those of previous studies suggesting high genetic control over the marginal effects of environment for the final bud set stages in boreal conifers (Perrin et al., 2017). The highly significant effect of the day of the year reflects the temporal aspect of phenology, which is mainly influenced by changes in temperature but also photoperiod (Li et al., 1993). Furthermore, large genetic variation in the timing of bud set has been observed among natural populations of white spruce in the northeastern part of its natural range, with southern populations exhibiting later bud set and, consequently, a longer growing season that resulted in greater growth (Li et al., 1993; Jaramillo-Correa et al., 2001). The same adaptive patterns were observed for black spruce in eastern Canada (Beaulieu et al., 2004; Perrin et al., 2017) and interior spruce in western Canada (Liepe et al., 2016).

Significant G×E interactions were recorded for the Desroberts and Baby seed sources, and for local seed sources. These interactions of northwestern and local seed sources suggest that bud set expression changes under warm conditions with longer growing seasons. This was observed through the increased likelihood probability reaching bud set stages 2 and 4 compared to more southern seed sources. Thus, northern seed sources could benefit from longer growing seasons under warming climate, allowing them to increase their growth through the extension of the growing season, while still maintaining a high level of frost tolerance.

### Survival rate of seed sources at test sites

The survival rate of seed sources at the northern Rousseau test site was above 75% for three seed sources: Desroberts, Falardeau (from the northwest and mid-latitude), and Fontbrune (from the south), while the three other seed sources (Baby from the western mid-latitude, Robidoux from the northeast, and Wendover from the south) showed a lower survival rate of around 60%. At the southern Wendover test site, most seed sources exhibited a high survival rate above 70%, except for the Baby seed source, which had a survival rate of only 50%. Notably, local seed sources showed approximately the same survival rate above 75% in both test sites. These results may underly adaptive genetic variation associated with local climates of seed source origins, given that many significant relationships have been observed between seed source variation, allelic variation at adaptive genes, and diverse factors related to climate in white spruce and other conifers in the region, with climatic variation following mainly a north-south latitudinal gradient in temperature and east-west longitudinal gradient in precipitation (Li et al., 1997; Beaulieu et al., 2004; Namroud et al., 2008; Prunier et al., 2011, 2013; Hornoy et al., 2015; Villeneuve et al., 2016; Depardieu et al., 2020, 2021). The survival rate was quite lower compared to previous studies involving the same seed sources and study area, which reported a 98% survival rate at the early stage after plantation (Villeneuve et al., 2016; Otis Prud’homme et al., 2018). Besides the increasing occurrence of drought episodes during the growing season in temperate and boreal ecozones at mid-latitudes (Depardieu et al., 2020), which could severely affect the ability of young white spruce trees to survive and grow given the limited development of their root systems (Soro et al., 2023), the decreased survival rate could also be explained, in part, by the increasing occurrence of early warm springs followed by late frosts. Indeed, these frosts are often happening while the new shoots have well initiated their elongation through an early start of the growing season, as was observed during spring 2021, with more intense effect in the south (Benomar et al., 2022). The proportion of damaged terminal shoots in this study confirmed this hypothesis by recording high damage rates exceeding 80% in the more meridional study site for all seed sources, compared to just over 60% in the more northern site.

### Frost damage and tolerance of seed sources in contrasted environments

The relative electrolytic conductivity (REC) and leaf damage index (LDI) of seed sources, measured across both test sites and under different freezing temperatures, showed significant effects of site, freezing temperature, their interaction, and the three-way interaction (site x seed source x freezing temperature), accounting for more than 85% of the total variance for each index. These two parameters were used as indirect methods in many previous studies to assess frost tolerance in various coniferous species, such as jack pine, black spruce, lodgepole pine, and white spruce (Lamhamedi et al., 2005, 2022; Man et al., 2021). Our results showed that most seed sources become vulnerable to frost damage under -12 °C when established in the meridional site. This level of vulnerability remains the same under -16 °C except for the mid-latitude Falardeau seed source and the meridional Wendover seed source. These two seed sources exhibited the highest levels of vulnerability to frost damage (Figure 8). However, all seed sources maintained high levels of frost tolerance when established in the northern site, even under -12 °C and -16 °C.

Although we detected a local adaptation effect among the different seed sources, our results primarily underline the importance of phenotypic plasticity of frost tolerance following the studied climatic gradient. Previous studies on ecophysiological traits of white spruce confirmed that the values of functional traits vary significantly between sites (Benomar et al., 2016). Also in Douglas fir, Schuch et al. (1989a, b) found that the cold hardiness of seed sources was mainly affected by plantation locations in the more meridional area of Oregon State, with those established higher north showing the highest frost tolerance and the earliest build-up of cold hardiness. Similarly, Repo et al. (2000) reported that northern seed sources of Scots pine showed an earlier onset of hardening and an increased build-up of frost tolerance. For white spruce, Lamhamedi et al. (2005) found that the development of frost tolerance is a progressive process, varying significantly over time (from the end of August to mid-October). The general pattern is that, during the end of summer and fall seasons, bud formation and the development of frost tolerance are closely linked to the decreasing photoperiod, accompanied by a drop in temperature (Bigras & D’Aoust, 1992; Li et al., 1993; Lamhamedi & Bernier, 1994; Johnsen & Skrøppa, 2000; Howe et al., 2003). In the light of our frost tolerance analysis, most seed sources planted in the north acquired frost tolerance under -16 °C earlier and showed no damage to tested samples. Among more southern seed sources, the southwestern Fontbrune seed source showed moderate frost tolerance when planted in the south compared to the Wendover seed source or the mid-latitude Falardeau seed source (same longitude as Wendover seed source). Thus, Fontbrune is likely to be a good candidate for northward transfer of white spruce under future warming climate conditions, where it will benefit from warmer conditions for growth and moderate frost tolerance. This hypothesis could be further confirmed through the sampling over many time points, particularly after the end of cold hardiness phenology, to better understand the timing of cold resistance acquisition at the end of the growing season.

### Growth performance of different seed sources in contrasted environments

The 7-year tree height showed highly significant effects of seed sources, test sites, and their interaction. The average height of trees of all seed sources combined was higher in the more meridional test site, highlighting the effect of the warmer climate conditions on growth productivity at this site. The Wendover southern seed source and Desroberts northern seed source showed similar performance in the more meridional test site. Our results are partially aligned with the findings of Benomar et al. (2016) and Otis Prud’homme et al. (2018) regarding the growth performance of the same white spruce seed sources in contrasting environments. They reported significant effects of the test site and the studied seed sources at their juvenile stage. However, no significant genotype x environment (G×E) interactions were reported, presumably because the trees were assessed at a very young age (3 to 4 years). However, Lu et al. (2014) reported significant G×E interactions for different white spruce seed sources’ growth performance in contrasted environments. Xie, (2003) has also reported a highly significant G×E interaction effect on 10-year height among 232 open-pollinated interior spruce families in north central interior British Columbia, where altitudinal variation among seed sources is much more pronounced than in eastern Canada. In our study, the G×E interaction implicating mostly the Fontbrune southern seed source could be explained by genetic adaptation to the local climate of origin, which is characterized by minimal and maximal daily extreme temperatures (T_min_ and T_max_), together with high values of precipitation variables (maximum 1-day precipitation and maximum 5 consecutive days of precipitation) (Supplementary Figures S5, S6). In fact, the climate profile of the Fontbrune seed source origin had similarities with the climate envelopes of both test sites. Combined with the moderate vulnerability of this seed source to frost damage (Figure 6), these adaptations could explain the important shift of the Fontbrune tree height when established in the south. Thus, the Fontbrune seed source would benefit more than others from longer growing seasons under future warming climate conditions.

### Dynamics of non-structural carbohydrates (NSC)

The sugar analysis demonstrated the great phenotypic plasticity of white spruce. Indeed, highly significant effects of the test site were recorded for contents in fructose, glucose, sucrose, and pinitol, suggesting strong environmental effects on non-structural carbohydrate content, which contributes to cold hardiness and thus to the build-up of frost tolerance (recall of a reference needed here). Fructose, glucose, and pinitol content decreased northwardly, reflecting their use during the growing season under warmer conditions, while sucrose increased northwardly, reflecting its role in the initiation of dormancy and cold resistance acquisition under cold conditions (Figure 8). The recorded correlations between the NSC contents reflect their interrelated metabolism. In fact, the ability of plants to synthesize and accumulate sucrose in leaves is mainly determined by the concerted action of three enzymes: sucrose degrading sucrose synthase (SS), acid invertase (AI), and sucrose accumulating sucrose phosphate synthase (SPS) (Rinne et al., 2015). The SS reactions are reversible, whereas AI reactions, which hydrolyze sucrose into glucose and fructose, are irreversible (Li et al., 2006). Simard et al. (2013) reported that sucrose was the least abundant sugar under high growth, which may explain its increase northward in our study. On the other hand, high concentrations of sugars, which play a role in cryoprotection and osmotic adjustments (Hoch & Korner, 2003), were described as a strategy to survive frequent biomass losses caused by extreme climatic events, particularly during the growing season (Sveinbjörnsson, 2000). Other studies reported high levels of sugars during the growing season in trees growing under cold conditions (Kagawa et al., 2006; Simard et al., 2013), which could explain the observed northward increase in sucrose content associated with cold resistance acquisition in our study.

Significant effects of the sampling date were detected for fructose, glucose, and sucrose contents, which indicate the presence of a dynamic metabolism in response to environmental changes. The interaction between test site and sampling date was significant for fructose and glucose contents, with an increasing trend over time in the southern site and a decreasing trend in the northern site. In contrast, sucrose content exhibited the opposite trend. These spatiotemporal interactions suggest the presence of dynamic plasticity in NSC compounds expression. The negative associations of fructose to the numbers of icing days and frost days and its positive association with the growing season length thus suggest that fructose is more related to growth and warm conditions (Figure 9A).

### Frost tolerance association with the averages of growth, the NSC compounds content, and the climate of seed source origins

The leaf damage index (LDI) showed a negative correlation with height and a positive correlation with fructose content. This suggests that the taller the tree, the less it is exposed to frost damage, while higher fructose levels are associated with increased frost damage. Indeed, Marquis et al. (2020) found that taller white spruce trees are less exposed to frost damage, given that freezing temperatures are more likely close to the ground (Benomar et al., 2022). Additionally, Simard et al. (2013) also found that fructose is related to high growth conditions and tends to be low in colder regions, as observed in our study for the northern Desroberts seed source (Figure 10). Furthermore, previous studies have shown that soluble sugars, including sucrose and pinitol, accumulate in leaf tissues to enhance freezing tolerance in pine trees (Strimbeck et al., 2008; Angelcheva et al., 2014).

Regarding the climate of seed source origins, their typical lengths of growing season, the minimum daily value of maximal temperature, the minimum daily value of minimal temperature, and the precipitation intensity index were positively correlated with the leaf damage index. This pattern was particularly conserved from northern to southern seed source location origins. Knowing that the longer the growing season of more meridional seed sources, the more exposed they could be exposed to frost damage. The minimum values of daily minimum and maximum temperature, characterized mainly by the intersection of both climatic southern and northern climatic envelopes, as described in Supplementary Figure 3. ???voir mon commentaire? The precipitation intensity index, which is an important factor affecting tree growth, was also positively correlated with the leaf damage index. Moreover, higher precipitation intensity was also correlated with the length of the growing season, which could explain in part the higher frost damage observed in more meridional seed sources.

On the other hand, negative associations of the leaf damage index were observed with the number of frost days, icing days, the maximum length of the wet spells, and temperature range. Indeed, our principal component analysis of bioclimate factors at test sites and seed source origins showed that frost days and icing days characterizing more northern seed origin locations were negatively correlated with summer days, precipitation intensity, and length of growing season (Supplementary Figure 3), which could explain the higher leaf damage values observed in the most meridional seed source (Wendover). These negatively correlated bioclimatic variables at the test sites are more likely to induce cold hardiness and dormancy in trees, thus decreasing their susceptibility to leaf damage. These associations would be mainly due to phenotypic plasticity, as observed in the correlation heatmaps with and without test site averages (Figure 9A & B). Our findings are thus aligned with the results of Sebastian-Azcona et al. (2018), indicating a significant relationship between cold hardiness (at -30 °C) and several climatic variables, including the length of the growing season, the coldest monthly temperature, winter temperatures, mean annual temperature, and first frost date.

The relative electrolytic conductivity (REC) of seed sources was positively correlated with the leaf damage index and fructose content. Knowing that low REC values reflect high frost tolerance levels (Lamhamedi et al., 2005, 2022), white spruce trees with lower REC would tend to have low frost damage when established in northern regions. In part, this pattern would be due to genetic adaptation of seed sources to their local environmental conditions. On the other hand, negative correlations were recorded for tree height and pinitol content.

Tall trees generally exhibited low REC values (Supplementary Figure 4), which could be explained in part by the reduced exposure to cold temperature near the soil surface but also, due to the plasticity of frost tolerance as highlighted in our freezing tests (Figure 6). As for the negative association between pinitol content and REC (Supplementary Figure 4), it would highlight the important role of pinitol in the acquisition of tolerance to cold temperatures, as previously reported (Strimbeck et al., 2008; Angelcheva et al., 2014).

### Implications for assisted migration

Our findings highlight the genetic, environmental, and genotype-by-environment (G×E) effects on growth and bud set phenology of the studied seed sources across climate-contrasting testing environments. Our results also showed significant relationships between frost tolerance, non-structural carbohydrate (NSC) content, tree growth, and the climate of both seed source origins and test sites representative of the regional climates of main reforestation activities. The identified relationships can be used to identify the best matches between seed sources and the target sites for intraspecific assisted migration favoring high productivity and enhanced tolerance to severe frosts. Taken together, these results should help improve the accuracy of seed transfer models across Quebec by integrating frost tolerance in seed source selection, and guide efforts to deployment of genetically improved white spruce so to maintain productivity and adaptability under future climatic conditions. Potentially further enhanced by integrating a desirability function into the seed transfer model.

## Conclusion and perspectives

The climate of seed source origins and the actual climate means at test sites showed significant correlations with the two frost tolerance indices, tree height, and NSC compounds. These relationships have allowed to map the complex relationships between growth, frost tolerance, and the roles of non-structural carbohydrate compounds in tree growth and cold resistance, in relation to site climate conditions and seed origins. The onset of bud set phenology was found to be similar for all seed sources in each test site but was earlier in the more meridional test site for all seed sources. The significant plasticity and dynamics of the non-structural carbohydrates and the identified spatio-temporal patterns suggested that dynamic changes are taking place at the end of the growing season and play a crucial role in cold hardiness and frost tolerance. This study highlighted adaptive genetic adaptation and the plasticity of white spruce seed sources, and it provides direct evidence that the northward transfer of southern seed sources in the southern Quebec conditions is not associated with a higher risk of frost sensitivity. Further studies investigating the dynamic aspects of cold hardening and non-structural carbohydrates from the first stage of bud set to the end of eco-dormancy would be key to expand the results here reported, and to determine whether the trends seen in the various ecophysiological processes studied here are amplified at the more large-scale distribution level of this widely distributed conifer species across the temperate and boreal forests of Canada.

## Data Availability Statement

The raw data and scripts supporting the results reported in this article will be made available by the authors upon request.

## Author Contributions

CA collected data, conducted statistical analyses, and drafted the manuscript.

LB collected data, reviewed and improved the manuscript.

MP and JG participated in obtaining funds, providing important support and access to the MRNF laboratory and plantation tests, and reviewed and improved the manuscript.

YB and JBe participated in obtaining funds and helped drafting the manuscript.

JBo helped plan and design the study, drafting the manuscript and funding of this project.

ML planned and designed the study, data collection, and drafting the manuscript and funding of this project.

All authors reviewed the manuscript and approved the final draft.

## Funding

Funding for this project was obtained from an NSERC Alliance grant (ALLRP 560992-20) to ML, JBo, YB, JBe, MP, and JG. The study sites used in the study are part of project number [112332078] conducted at the Direction de la recherche forestière (Ministère des Ressources naturelles et des Forêts, Québec, Canada) and led by Martin Perron.

## Conflict of Interest

The authors declare that the research was conducted in the absence of any commercial or financial relationships that could be construed as a potential conflict of interest.

## Acknowledgments

We thank the Direction de la recherche forestière of the Ministère des Ressources naturelles et des Forêts du Québec for access to their laboratory and the study sites, and for the establishment and maintenance of the plantation tests used in this study.

---

**Supplementary Figure S1.**
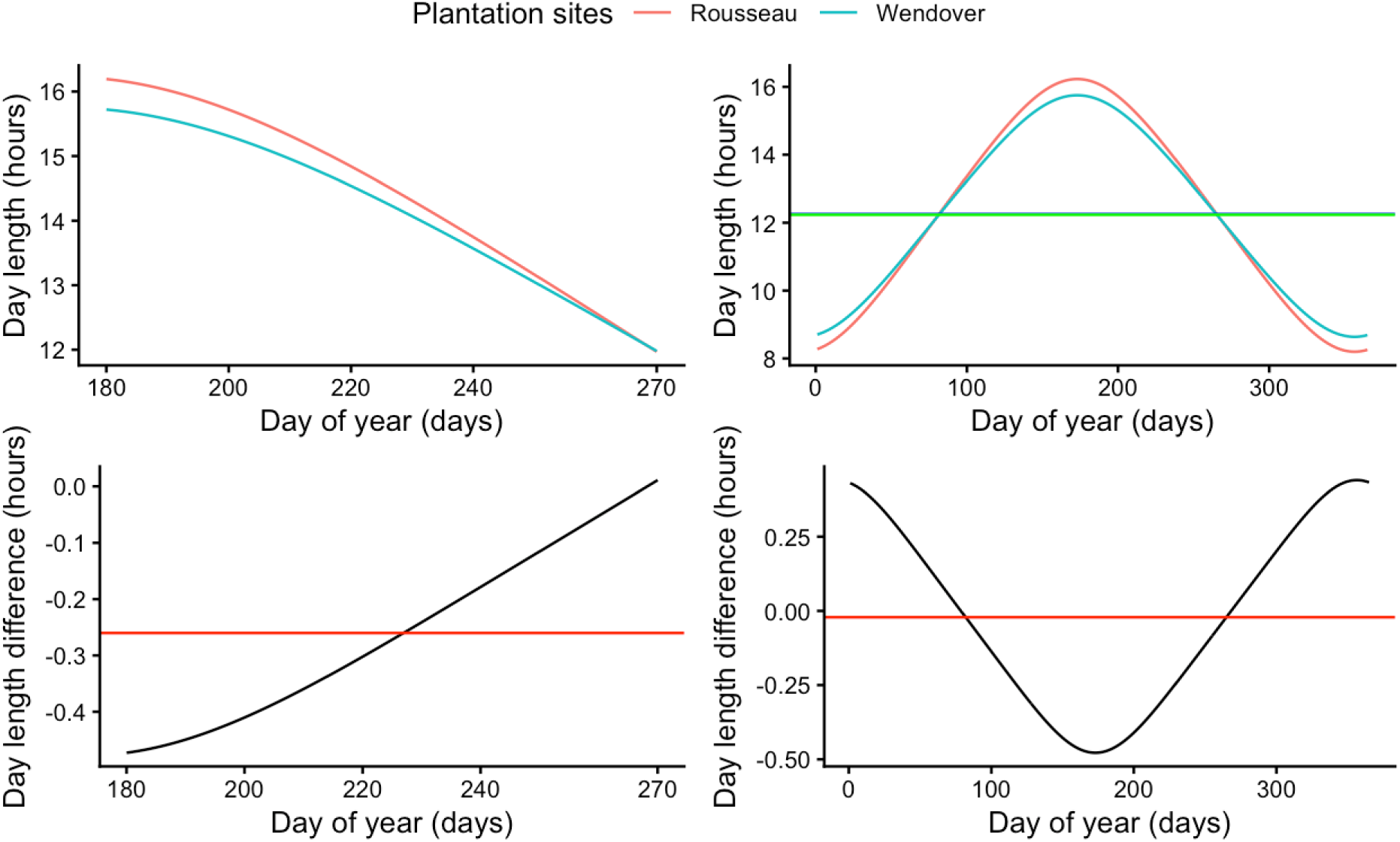
Day length distribution over a year for both test sites and the observed differences between them. The distribution was also recorded from the first day of budset to the last day of the year. The red and green horizontal lines represent the average day length and average day length difference between site over the corresponding period.

**Supplementary Figure S2.**
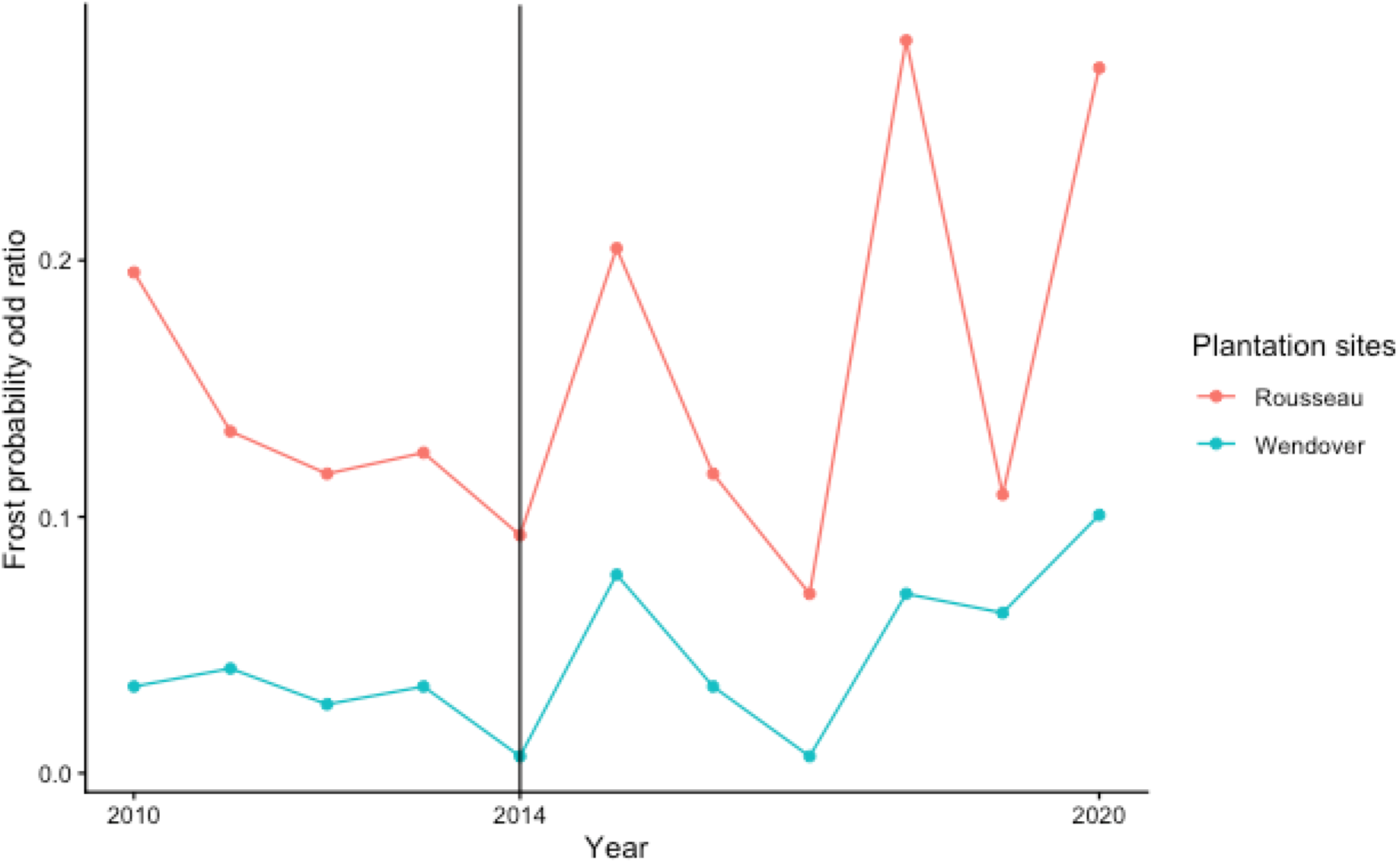
Annual odd ratio of frost probability variation recorded during the growing season over a 30-year period for the northern Rousseau and the southern Wendover test sites. The vertical line represents the planting year (2014), where the climate heterogeneity starts.

**Supplementary Figure S3:**
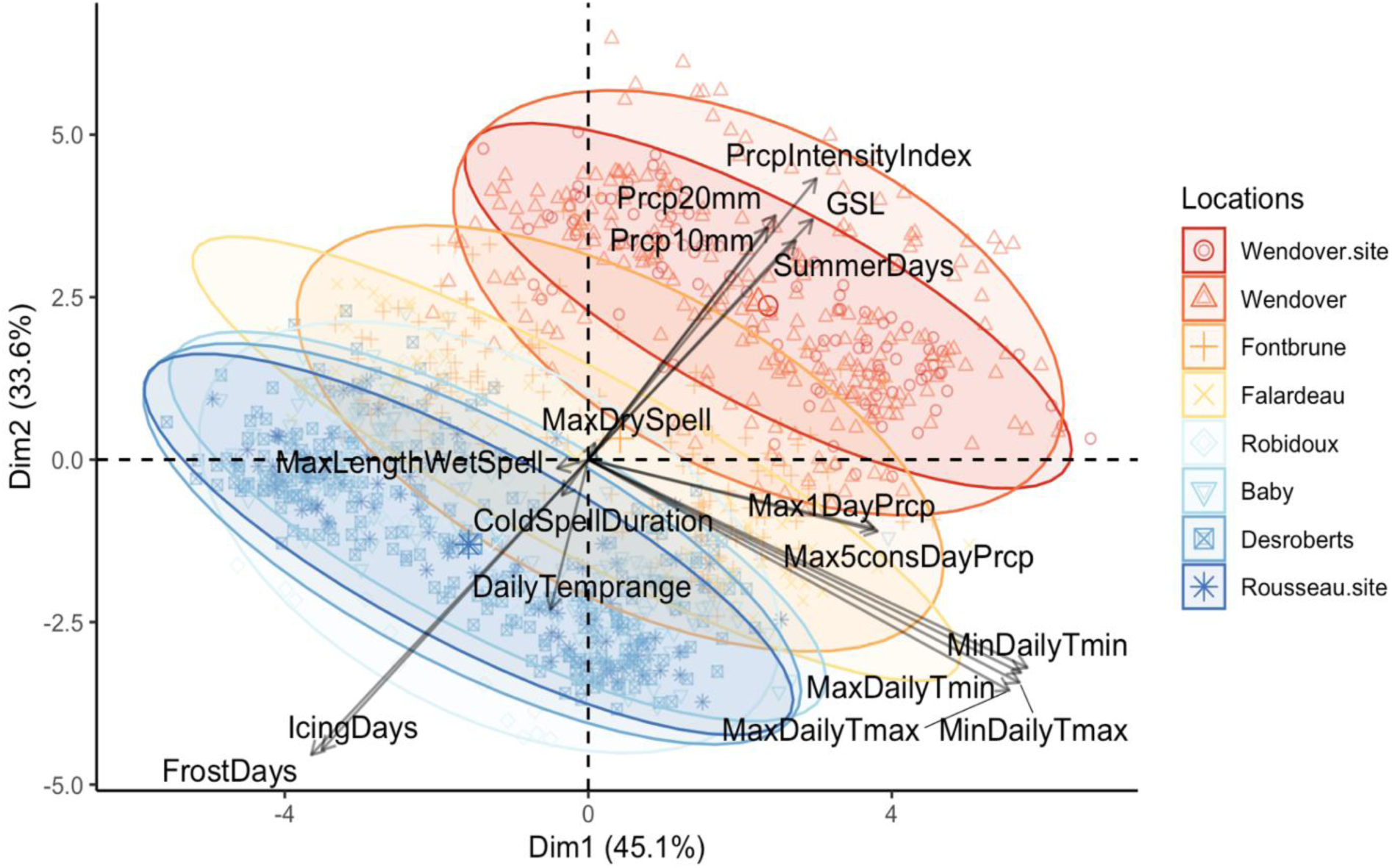
Biplot of principal component analysis (PCA) of bioclimatic variables from seed source origins and test sites. Locations in the legend are ordered geographically from south (Wendover) to north (Desroberts seed source and Rousseau test site).

**Supplementary Figure S4:**
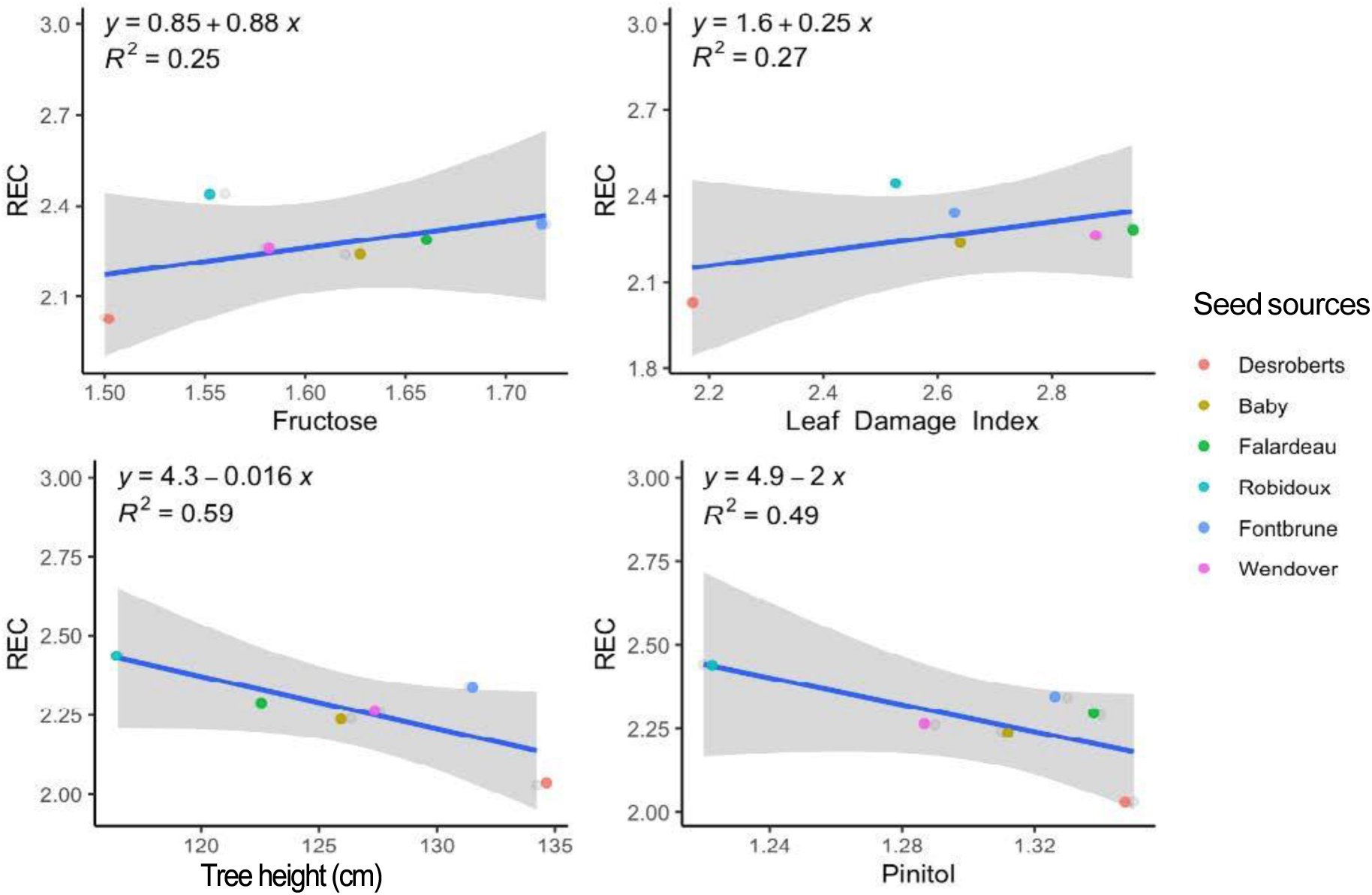
Simple linear regressions of mean relative electrolytic conductivity (REC) and confidence intervals versus non-structural carbohydrates content and tree height of seed sources.

**Supplementary Figure S5:**
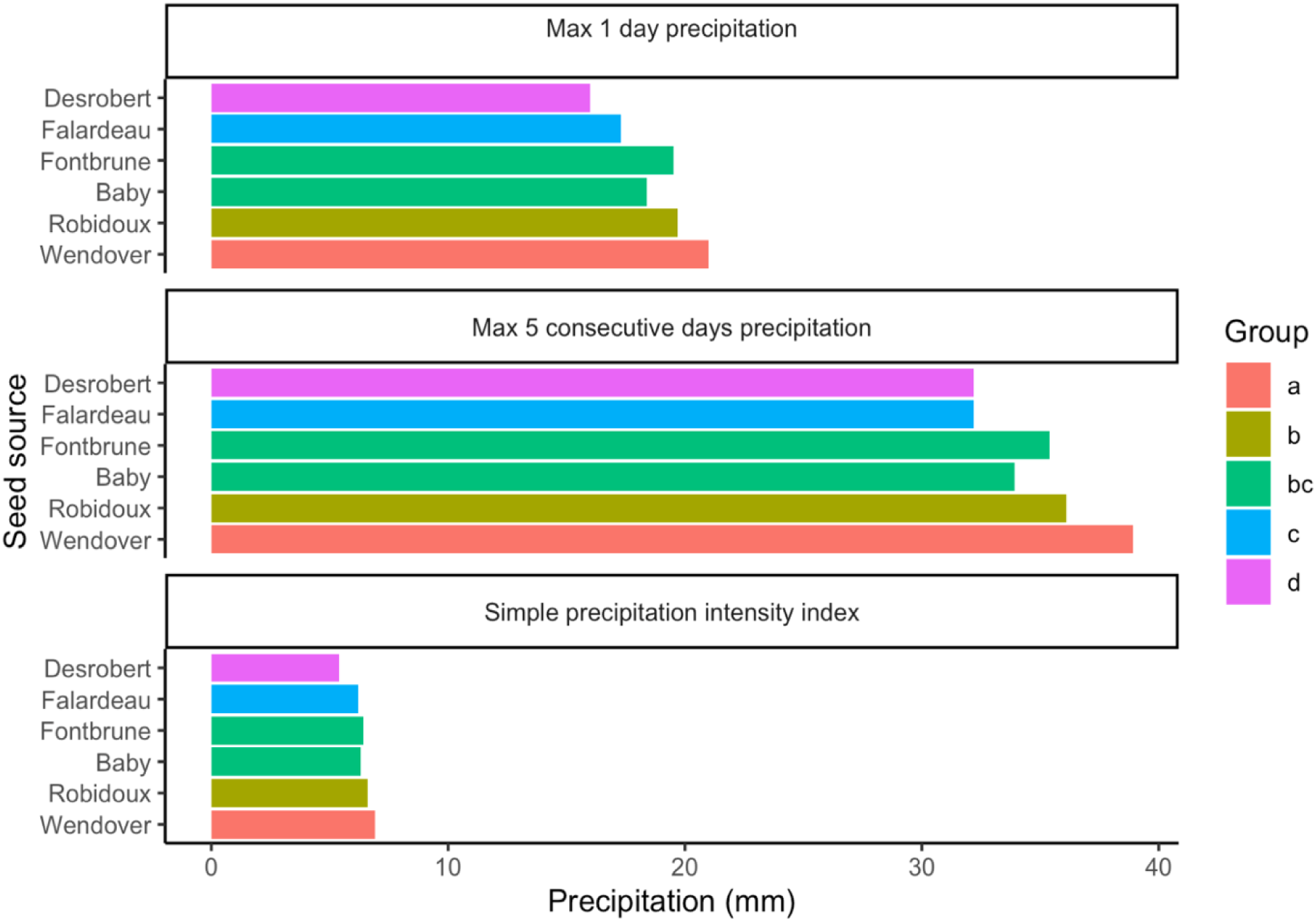
Distribution of the annual means of precipitation indices of seed sources calculated for their climate of origin (1940-1970). The groups represent the results of Tukey’s honest significant tests. https://www.climdex.org/learn/indices/.

**Supplementary Figure S6:**
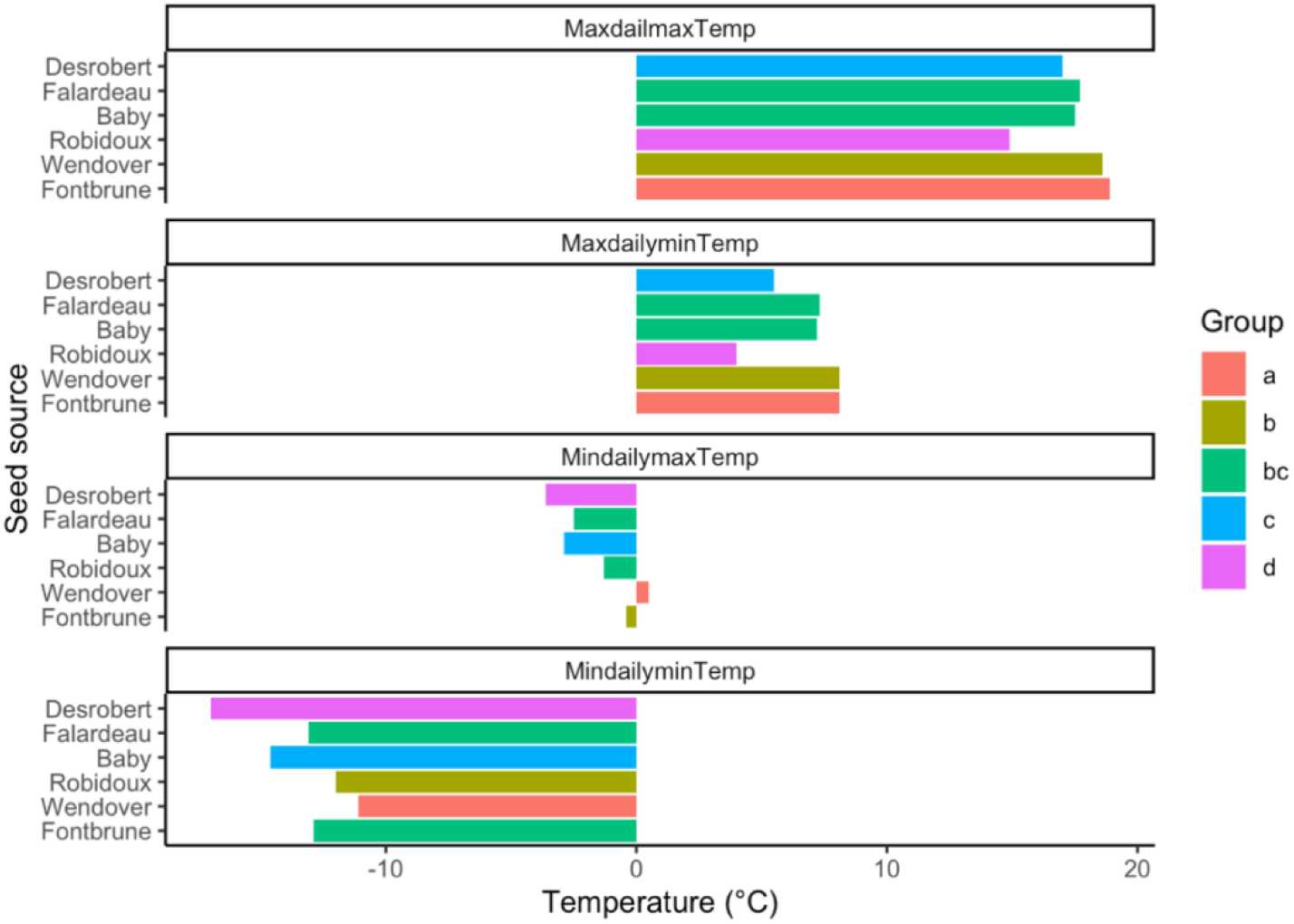
Distribution of *annual mean temperature extreme* indices for seed sources, calculated on their climate of origin (1940-1970). The groups represent the results of Tukey’s Honest Significant Difference (HSD) test. *Indices are defined according to ClimDex* https://www.climdex.org/learn/indices/.

**Supplementary Table S1:**
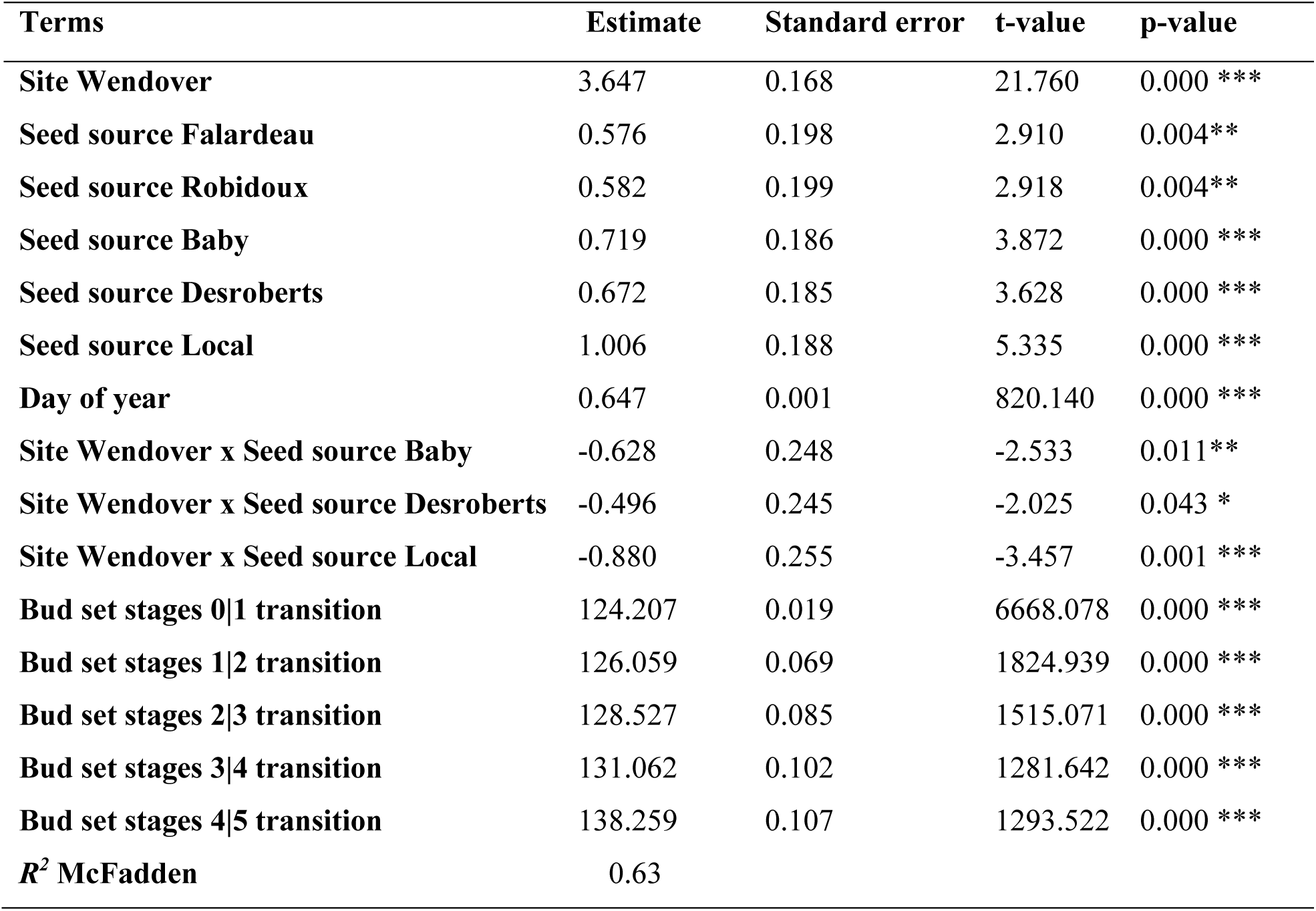
Summary statistics of ordinal logistic regression of phenology under test sites, seed sources, their interaction, and Julian day of the year effects. Only significant effects are reported. Note: The estimates for the Wendover test site were calculated with the Rousseau test site being the reference. For seed sources estimates, the Wendover seed source was the reference. The bud set stages transitions are expressed as follows: Stage n | Stage n+1 (n starting from 0). The *R^2^* McFadden reflects the total variance explained by the model. It is equivalent to *R^2^* coefficient for linear models.

## Notes

### Competing Interest Statement

The authors have declared no competing interest.

